# Single-cell Ca^2+^ parameter inference reveals how transcriptional states inform dynamic cell responses

**DOI:** 10.1101/2022.05.18.492557

**Authors:** Xiaojun Wu, Roy Wollman, Adam L. MacLean

**Affiliations:** Department of Quantitative and Computational Biology, University of Southern California, 1050 Childs Way, Los Angeles, CA 90089, USA; Departments of Integrative Biology and Physiology and Chemistry and Biochemistry, University of California Los Angeles. Los Angeles, CA

## Abstract

Single-cell genomic technologies offer vast new resources with which to study cells, but their potential to inform parameter inference of cell dynamics has yet to be fully realized. Here we develop methods for Bayesian parameter inference with data that jointly measure gene expression and Ca^2+^ dynamics in single cells. We propose to share information between cells via transfer learning: for a sequence of cells, the posterior distribution of one cell is used to inform the prior distribution of the next. In application to intracellular Ca^2+^ signaling dynamics, we fit the parameters of a dynamical model for thousands of cells with variable single-cell responses. We show that transfer learning accelerates inference with sequences of cells regardless of how the cells are ordered. However, only by ordering cells based on their transcriptional similarity can we distinguish Ca^2+^ dynamic profiles and associated marker genes from the posterior distributions. Inference results reveal complex and competing sources of cell heterogeneity: parameter covariation can diverge between the intracellular and intercellular contexts. Overall, we discuss the extent to which single-cell parameter inference informed by transcriptional similarity can quantify relationships between gene expression states and signaling dynamics in single cells.

## 1 Introduction

Models in systems biology span systems from the scale of protein/DNA interactions to cellular, organ, and whole organism phenotypes. Their assumptions and validity are assessed through their ability to describe biological observations, often accomplished by simulating models and fitting them to data [1, 2, 3, 4]. Under the framework of Bayesian parameter inference and model selection, the available data is used along with prior knowledge to infer a posterior parameter distribution for the model [5]. The posterior distribution characterizes the most likely parameter values to give rise to the data as well as the uncertainty that we have regarding those parameters. Thus, parameter inference provides a map from the dynamic phenotypes that we observe in experiments to the parameters of a mathematical model.

Single-cell genomics technologies have revealed a wealth of information about the states of single cells that was not previously accessible [6]. This ought to assist with the characterization of dynamic phenotypes. However, it is much less clear how to draw maps between dynamic phenotypes of the cell and single-cell states as quantified via genomic measurements. The challenge in part lies in the combinatorial complexity: even if a small fraction of genes contain information regarding the phenotype of interest, say a few hundred, this is more than enough to characterize any feasible number of states of an arbitrarily complex dynamical process.

This leads us to a central question: can the integration of single-cell gene expression data into a framework for parameter inference improve our understanding of the cellular phenotypes of interest? Here, various sources of transcriptional noise must be taken into account [7, 8, 9]; which we propose to address by taking a global view and comparing cells first by their similarity across many genes, and, after inference, by their similarity in posterior parameter distributions. Our previously work provides the ideal data for this approach: we jointly measured dynamics and gene expression in the same single cells [10]. Here, we apply our new parameter inference framework to study Ca^2+^ signaling dynamics and signal transduction in response to adenosine triphosphate (ATP) in human mammary epithelial (MCF10A) cells.

Ca^2+^ signaling regulates a host of cellular responses in epithelial cells, from death and division to migration and molecular secretion, as well as collective behaviors, such as organogenesis and wound healing [11, 12, 13]. In response to ATP binding to purinergic receptors, a signaling cascade is initiated whereby phospholipase C (PLC) is activated and in turn hydrolyzes phosphatidylinositol 4,5-bisphosphate (PIP2), producing inositol 1,4,5-trisphosphate (IP3) and diacylglycerol (DAG). The endoplasmic reticulum (ER) responds to IP3 by the activation of Ca^2+^ channels: the subsequent release of calcium from the ER into the cytosol produces a spiked calcium response. To complete the cycle and return cytosolic calcium levels to steady state, the sarco/ER Ca^2+^-ATPase (SERCA) channel pumps the Ca^2+^ from the cytosol back into the ER [14, 15]. Ca^2+^ signaling is highly conserved, regulating cell phenotypic responses across mammals, fish, and flies [12, 16, 17] as well as in prokaryotes [18]. Since the Ca^2+^ response to ATP occurs quickly in epithelial cells: on a timescale that is almost certainly faster than gene transcription, we work under the assumption that the transcriptional state of the cell does not change in the duration of the experiment.

Our ability to measure gene expression in thousands of single cells has not only led to new discoveries but has also fundamentally changed how we identify and characterize cell states [19]. Technologies used to quantify gene expression in single cells include sequencing and fluorescent imaging. The latter permits the measurement of hundreds of genes in spatially-resolved populations of single cells. Small molecule fluorescence in situ hybridization (smFISH) can be multiplexed to achieve this high resolution by protocols such as MERFISH [20] and seqFISH [21]. Moreover, by coupling multiplexed smFISH with fluorescent imaging of Ca^2+^ dynamics using a GFP reporter in MCF10A cells, we are able to jointly capture the dynamic cell responses and the single-cell gene expression in the same single cells [10]. These data offer new potential to study the relationships between transcriptional states of cells and the dynamic phenotypes these may produce.

Models of gene regulatory networks and cellular signaling pathways described by ordinary differential equations (ODEs) capture the interactions between gene transcripts, proteins, or other molecular species and their impact on cellular dynamics. Well-established dynamical systems theory offers a range of tools with which to analyze transient and equilibrium behavior of ODE models [22]; it remains an open question whether or not such it is appropriate to make equilibrium assumptions of living cells [23]. Constraining dynamic models of cellular/molecular processes with single-cell data via inference offers much potential to gain new insight into dynamics, albeit coming with many challenges given, among other things, the complex sources of noise in these data and the lack of explicit temporal information in (“snapshot”) datasets gathered at one time point [24]. Parameter inference has provided insight into clonal relationships of single cells [25, 26] and stem cell differentiation/cell state transitions [27, 28]. Inference methods have also been applied to single-cell data for the discovery of new properties of single-cell oscillations [29, 30] and cell-cell variability [31, 32, 33], as well as to study cell-cell communication [34]. New methods to infer the parameters of models of stochastic gene expression provide means to study single-cell dynamics in greater depth [35, 36].

Here, we model Ca^2+^ dynamics via ODEs based on previous work [37, 38]. We develop a parameter inference framework to fit Ca^2+^ response dynamics in many single cells. We perform inference of multiple cells sequentially, through the construction of “cell chains.” A cell chain is an ordering of cells, which can be random or directed by some measure, e.g. by similarity of gene expression or of Ca^2+^ dynamic response. Given a cell chain, we propose to infer the parameters of the Ca^2+^ ODE model in a single cell via a transfer of information from its cell predecessor in the chain. We achieve this by setting the prior of the current cell in the chain informed by the posterior of its predecessor. We will use this framework to assess the extent to which transcriptional cell states inform dynamic cell responses.

In the next section we present the model and the methods implemented for parameter inference using Hamiltonian Monte Carlo in Stan [39]. We go on to study the results of inference: we discover that priors informed by cell predecessors accelerate parameter inference, but that cell chains with randomly sampled predecessors perform as well as those with transcriptional similarity-informed predecessors. Analysis of hundreds of fitted single cells reveals that cell-intrinsic vs. cell-extrinsic posterior parameter relationships can differ widely, indicative of fundamentally different sources of underlying variability. By perturbing the posterior distributions, we assess model parameter sensitivities in light of Ca^2+^ dynamics. We also find that variability in single-cell gene expression is associated with variability in posterior parameter distributions, both for individual gene-parameter pairs and globally, via principal component analysis. We go on to cluster cells by their posterior parameter distributions, and discover that only for gene expression-based cell chains are there clear relationships between gene expression states and dynamic cell phenotypes.

## 2 Materials and Methods

### 2.1 A model of Ca^2+^ dynamics in response to ATP

We model Ca^2+^ signaling pathway responses in MCF10A human epithelial cells using nonlinear ordinary differential equations (ODEs), as previously developed [37, 38]. The model consists of four state variables: phospholipase C (PLC), inositol 1,4,5-trisphosphate (IP3), the fraction of IP3-activated receptor (*h*), and cytoplasmic Ca^2+^. The four variables are associated with a system four nonlinear ODEs describing the rates of change of the Ca^2+^ pathway species following ATP stimulation, to characterize dynamic responses in MCF10A cells. The equations are given by:

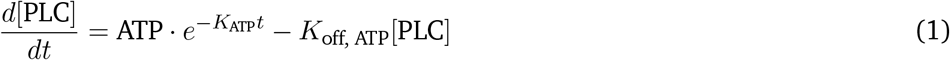

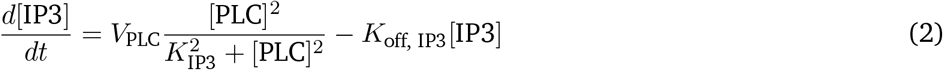

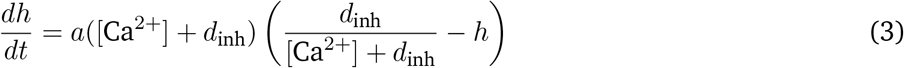

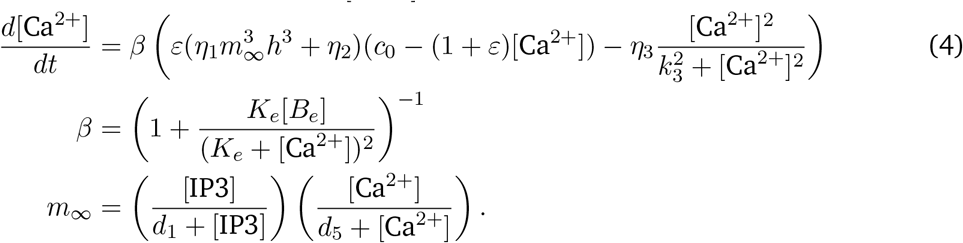

The equations describe a chain of responses following ATP binding to purinergic receptors: the activations of PLC, IP3, the IP_3_R channel on the surface of the ER, and finally the release of Ca^2+^ from the ER into the cytoplasm [38]. Ca^2+^ may also enter the ER through the IP_3_R channel and the SERCA pump [38]. Our model differs from Yao et al. [38] in that we combine the product of two parameters in the previous model, *K*_on, ATP_ and ATP, into a single parameter, ATP. This reduction of the model parameter space removed the redundancy that would otherwise exist in the distributions of *K*_on, ATP_ and ATP. A description of each of the parameters in the model is given in Table 1, where reference values for each of the model parameters are found in Lemon et al. [37] and Yao et al. [38].

**Table 1:** Definition and description of the ODE model parameters. Prior distributions are derived from [37, 40, 38].

| Name | Description | Prior distribution | Unit |
| --- | --- | --- | --- |
| ATP | Concentration of ATP that activates PLC | $\mathcal{N}(5, 4)$ | $\text{s}^{-1}$ |
| $K_{\text{ATP}}$ | ATP decay rate | $\mathcal{N}(0.0083, 0.0025)$ | $\text{s}^{-1}$ |
| $K_{\text{off, ATP}}$ | PLC degradation rate | $\mathcal{N}(1.25, 1)$ | $\text{s}^{-1}$ |
| $V_{\text{PLC}}$ | Maximum velocity for IP3 generation | $\mathcal{N}(1, 1)$ | $\mu\text{M}\cdot\text{s}^{-1}$ |
| $K_{\text{IP3}}$ | Equilibrium constant for IP3 generation through PLC | $\mathcal{N}(0.5, 0.01)$ | $\mu\text{M}$ |
| $K_{\text{off, IP3}}$ | IP3 degradation rate | $\mathcal{N}(1.25, 1)$ | $\text{s}^{-1}$ |
| $a$ | Time constant of IP3 channel | $\mathcal{N}(1, 1)$ | $\text{s}^{-1}$ |
| $d_{\text{inh}}$ | Dissociation constant for IP3 channel calcium inhibiting subunit | $\mathcal{N}(0.4, 0.01)$ | $\mu\text{M}$ |
| $K_e$ | Dissociation constant for calcium buffer | $\mathcal{N}(10, 4)$ | $\mu\text{M}$ |
| $B_e$ | Concentration of calcium buffer | $\mathcal{N}(150, 25)$ | $\mu\text{M}$ |
| $d_1$ | Dissociation constant for IP3 channel IP3 activating subunit | $\mathcal{N}(0.13, 0.01)$ | $\mu\text{M}$ |
| $d_5$ | Dissociation constant for IP3 channel calcium activating subunit | $\mathcal{N}(0.0823, 0.01)$ | $\mu\text{M}$ |
| $\epsilon$ | ER to cytosolic volume | $\mathcal{N}(0.185, 0.01)$ | – |
| $\eta_1$ | IP3 channel permeability constant | $\mathcal{N}(575, 625)$ | $\text{s}^{-1}$ |
| $\eta_2$ | ER leak permeability constant | $\mathcal{N}(5.2, 1)$ | $\text{s}^{-1}$ |
| $\eta_3$ | $\text{Ca}^{2+}$ pump permeability constant | $\mathcal{N}(45, 25)$ | $\text{s}^{-1}$ |
| $c_0$ | Concentration of free $\text{Ca}^{2+}$ in the ER | $\mathcal{N}(4, 1)$ | $\mu\text{M}$ |
| $k_3$ | SERCA pump dissociation constant | $\mathcal{N}(0.4, 0.01)$ | $\mu\text{M}$ |

### 2.2 Data collection and preprocessing

The data consist of a joint assay measuring Ca^2+^ dynamics and gene expression via multiplexed error-robust fluorescence *in situ* hybridization (MERFISH) [20]. Ca^2+^ dynamics in a total of 5128 human MCF10A cells are measured via imaging for 1000 seconds (ATP stimulation at 200 seconds) using a GCaMP5 biosensor. Immediately following this step, 336 genes are measured by MERFISH [10]. The Ca^2+^ trajectories are smoothed using a moving average filter with a twenty-second window size (Figure S1). After smoothing, data points occurring before ATP stimulation are removed. Data points for each Ca^2+^ trajectory after *t*=300 are downsampled by a factor of 10; the trajectories are at or close to steady state by this time. After removing the data for the first 200 seconds and downsampling for the last 700 seconds, each processed trajectory consists of Ca^2+^ response data on 171 time points (*t* = 200, 201, …, 298, 299, 300, 310, 320, …, 1000). All numerical experiments in this work will evaluate Ca^2+^ response on those same 171 time points. Single-cell gene expression data is collected using MERFISH after the Ca^2+^ imaging as previously described [10, 20].

### 2.3 Generating cell chains via cell-cell similarity

Cell-cell similarity is quantified via single-cell transcriptional states, i.e. by comparing 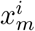 and 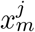, the expression of *m* genes in cells *i* and *j*. We obtain a symmetric cell-cell similarity matrix, *W*, from the log-transformed MERFISH expression data via optimization in SoptSC [41]: entries *W*_*i,j*_ denote the similarity between cells *i* and *j*. To create a chain of cells linked through their similarity in gene expression space, we:

1. Construct a graph *G* = (*V, E*); each node is a cell and an edge is placed between two cells if they have a similarity score above zero;
2. For a choice of initial (root) cell, traverse *G* and record the order of cells traversed.

Ideally, each cell would be visited exactly once, however this amounts to finding a Hamiltonian path in *G*, an NP-complete problem. Therefore, as a heuristic solution we use a depth-first search (DFS), which can be completed in linear time. From the current node, randomly select an unvisited neighbor node and set this as the next current node, recording it once visited (pre-order DFS). If the current node has no unvisited neighbors, backtracks until a node with unvisited neighbors is found. When there is no unvisited node left, every node in the graph has been visited exactly once. Given cases where the similarity matrix is sparse (as we have here), the DFS generates a tree that is very close to a straight path.

### 2.4 Bayesian parameter inference with posterior-informed prior distributions

We seek to infer dynamic model parameters in single cells, informed by cell-cell similarity via the position of a cell in a cell chain. We use the Markov chain Monte Carlo (MCMC) implementation: Hamiltonian Monte Carlo (HMC) and the No-U-Turn Sampler (NUTS) in Stan [42, 39]. HMC improves upon the efficiency of other MCMC algorithms by treating the sampling process as a physical system and employing conservation laws [43]. From an initial distribution, the algorithm proceeds through intermediate phase of sampling (warmup) until (one hopes) convergence to the stationary distribution. During warmup, NUTS adjusts the HMC hyperparameters automatically [42].

The prior distribution over parameters is a multivariate normal distribution, with dimensions *θ*_*j*_, *j* = 1, …, *m*, where *m* is the number of parameters. This choice of prior makes it straightforward to pass information from the inferred posterior distribution of one cell to the next cell in line to be sampled, which will be described in Section 2.5. Let *f* be a numerical solution of the ODE model, and *y*_0_ be the initial condition. Then, in each single cell, the Ca^2+^ response to ATP is generated by the following process:

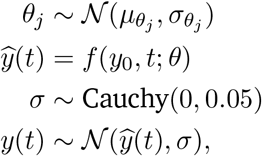

where we truncate the prior so that each *θ*_*i*_ is bounded by 0 from below. The Cauchy distribution is chosen to generate the noise for observed Ca^2+^ response as it contains greater probability mass in its tails, thus encouraging NUTS to explore extreme values of the parameter space more frequently.

For the first cell in a chain, we use a relatively uninformative prior, the “Lemon” prior (Table 1), derived from parameter value estimates in previous work [37, 40, 38]. For the *i*^th^ cell in a chain (*i* > 1), the prior distribution is constructed from the posterior distribution of the (*i* − 1)^th^ cell (Section 2.5). For each cell, NUTS is run for four independent chains with the same initialization. To simulate 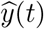 during sampling, we use the implementation of fourth and fifth order Runge-Kutta in Stan [39]. For each output trajectory *y*, its error is the Euclidean distance between *y* and data *y*\* for all 171 data points:

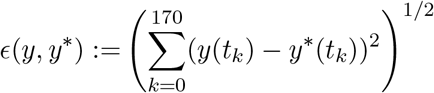

The error of a posterior sample for a cell is the mean error of trajectories simulated from all draws in the sample:

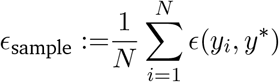

where *N* is the sample size and *y*_*i*_ is the output trajectory from the *i*^th^ draw in the sample.

Convergence of NUTS chains is evaluated using the 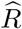 statistic: the ratio of between-chain variance to within-chain variance [39, 44]. A typical heuristic used is 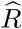 between 0.9 and 1.1 indicates that for this set of chains the stationary distributions reached for a given parameter are well-mixed. There are two caveats on our use of 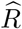 in practice:

1. For our model, we observe that well-fit (i.e. not overfit) Ca^2+^ trajectories did not require 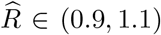 for all parameters. Thus we assess 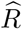 only for the log posterior, using a more tolerant upper bound of 4.0.
2. There are cases where one chain diverges but 3/4 are well-mixed. In such cases, we choose to retain the three well-mixed chains as a sufficiently successful run. Thus if 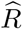 is above the threshold, before discarding the run, we compute 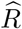 for all three-wise combinations of chains, and retain the run if there exist three well-mixed chains.

### 2.5 Constructing and constraining prior distributions

We construct the prior distribution of the *i*^th^ cell from the posterior of the (*i* − 1)^th^ cell. The prior mean for each parameter *θ*_*j*_ for the *i*^th^ cell is set to 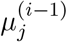, the posterior mean of *θ*_*j*_ from the (*i* − 1)^th^ cell. The variance of the prior for *θ*_*j*_ is derived from 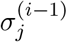, the posterior variance of *θ*_*j*_ from the (*i*− 1)^th^ cell. To a) sufficiently explore the parameter space, and b) prevent instabilities (rapid growth or decline) in marginal parameter posterior values along the cell chain, we scale each 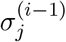 by a factor of 1.5 and clip the scaled value to be between 0.001 and 5. The scaled and clipped value is then set as the prior variance for *θ*_*j*_ for the *i*^th^ cell.

### 2.6 Dimensionality reduction and sensitivity analyses

To compare posterior samples from different cells, we use principal component analysis (PCA). Posterior samples are projected onto a subspace by first choosing a cell (the focal cell) and normalizing the posterior samples from other cells against the focal cell, either by min-max or *z*-score normalization. Min-max normalization transforms a vector *x* to 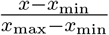, where *x*_min_ is the *x*_max_ minimum and the maximum of *x. z*-score normalization transforms *x* to 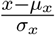, where *μ*_*x*_ is the mean and *σ*_*x*_ is the standard deviation of *x*. Normalizing to the focal cell amounts to setting *x*_min_, *x*_max_, *μ*_*x*_, *σ*_*x*_ to be the values corresponding to the focal cell for all cells normalized. We perform PCA (implemented by scikit-learn 0.24 [45]) on the normalized focal cell posterior samples and project them into the subspace spanned by the first two principal components. The normalized samples from all other cells are projected onto the PC1-PC2 subspace of the focal cell.

We develop methods for within-posterior sensitivity analysis to assess how perturbations of model parameters within the bounds of the posterior distribution affect Ca^2+^ responses. Given 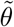, the posterior distribution of a cell, each parameter *θ*_*j*_ is perturbed to two extreme values: 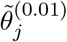, the 0.01-quantile of 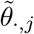, and 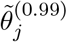, the 0.99-quantile of 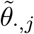. Nine “evenly spaced” samples are drawn from the posterior range of 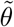 for the parameter of interest, 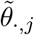: the *k* draw corresponds to a sample 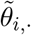 such that 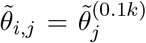, the 0.1*k*-quantile of 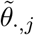. For each draw 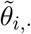, we replace 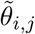 by either or 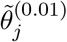 or 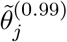 and then simulate a Ca^2+^ response. The mean Euclidean distances between trajectories simulated from the evenly spaced samples and the perturbed samples are used to quantify the sensitivity of each parameter perturbation.

### 2.7 Correlation analysis and cell clustering of MERFISH data

Correlations between single-cell gene expression values and posterior parameters from the Ca^2+^ pathway model are determined for variable genes. We calculate the *z*-scores of posterior means for each parameter of a cell sampled from a population, and remove that cell if any of its parameters has a posterior mean *z*-score smaller than −3.0 or greater than 3.0. PCA is performed on log-normalized gene expression of remaining cells using scikit-learn 0.24 [45], which yields a loadings matrix *A* such that *A*_*i,j*_ represents the “contribution” of gene *i* to component *j*. We designate gene *i* as variable if *A*_*i,j*_ is ranked top 10 or bottom 10 in the *j*^th^ column of *A* for any *j* ≤ 10. For each variable gene, we calculate the Pearson correlation between its log-normalized expression value and the posterior means of individual model parameters. Gene-parameter pairs are ranked by their absolute Pearson correlations and the top 30 are selected for analysis. Gene-parameter pair relationships are quantified by linear regression using a Huber loss, which is more robust to outliers than mean squared error.

To cluster cells using their single-cell gene expression, raw count matrices are normalized, log-transformed, and scaled to zero mean and unit variance before clustering using the Leiden algorithm at 0.5 resolution [46], implemented in Scanpy 1.8 [47]. Marker genes for each cluster are determined by a t-test.

### 2.8 Clustering of cell posterior parameter distributions

Cells are clustered according to their posterior distributions. For each parameter, the posterior means for each cell are computed and scaled to [0, 1]. The distance between two cells is defined as the *m*-dimensional Euclidean distance between their posterior means (where *m* is the number of parameters). Given distances calculated between all pairs of cells, agglomerative clustering with Ward linkage is performed using SciPy 1.7 [48]. Marker genes for each cluster identified are determined using a t-test.

## 3 Results

### 3.1 Single-cell priors informed by cell predecessors enable computationally efficient parameter inference

To study the dynamic Ca^2+^ responses of cells to ATP stimulation, we fit the ODE model (Eqns. (1–4)) to data in single cells using Bayesian parameter inference (Figure 1A). Only those MCF10A cells classified as “responders” to ATP were included — cells with very low overall responses (less than 1.8 Ca^2+^ peak height) were filtered out. To assess whether cell chains improve inference, we performed parameter inference of the ODE model in single cells fit either individually, each from the same prior (we used the “Lemon” prior (Table 1)), or fit via the construction of a cell chain. In a cell chain there is a transfer of information, whereby the posterior parameter distribution of one cell informs the prior distribution of the next cell in the chain. The first cell in the chain was fit using the Lemon prior. We are primarily interested in cell chains constructed using transcriptional similarity: we constructed cell chains based on a single-cell gene expression similarity metric and compared them with alternatives (see Methods, Table S1). We studied the effects of different choices of *g*, where *π*_*i*+1_ = *g*(*p*_*i*_); *p*_*i*_ is the posterior distribution for cell *i* and *π*_*i*+1_ is the prior distribution for the following cell. We found that transformations via scaling and clipping were necessary to sufficiently explore the parameter space for each cell while maintaining stable marginal posterior distributions along a cell chain (Section S1.1, Figure S2). We tested various numerical methods to solve the ODE system (stiff and non-stiff), and found that we could simulate Ca^2+^ responses sufficiently well using a non-stiff solver (Figure S3), so for inference runs with hundreds of single cells we proceeded to use a non-stiff solver.

**Figure 1:**
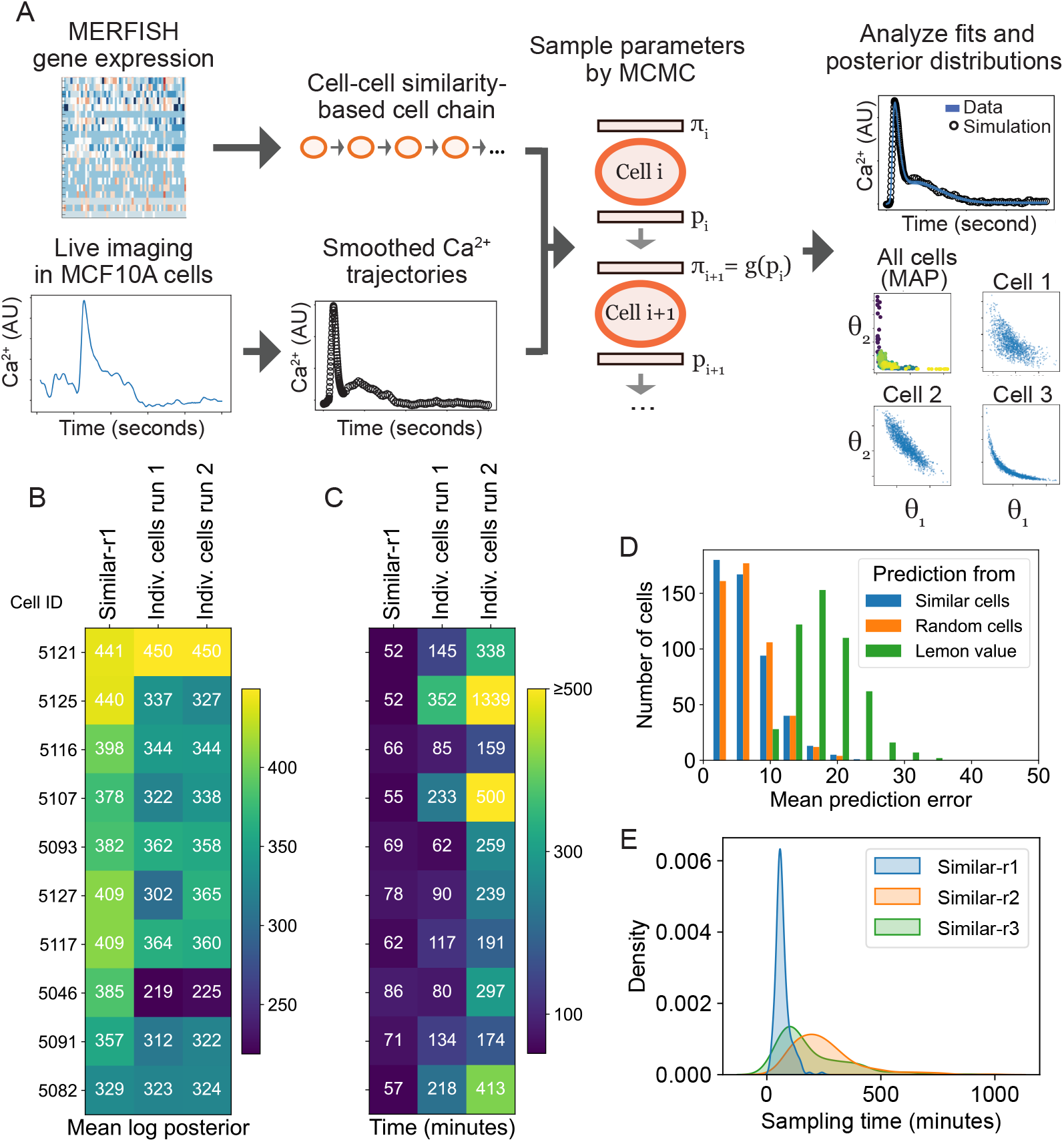
Cell chains improve performance of single-cell parameter inference. **A**: Workflow for single-cell parameter inference along a cell chain. *π*_*i*_ and *p*_*i*_ respectively denote the prior and posterior distributions of cell *i*. MAP: maximum a posteriori value. **B–C**: Comparison of inference for gene expression similarity-based cell chain (*Similar-r1*; first column) vs. cells fit individually, i.e. without a cell chain (second two columns). Output metrics are the mean log posterior, where higher values denote better fits; and the sampling times. Each row represents a cell. For *Similar-r1*, one cell was sampled from a Lemon prior before Cell 5121 to inform the prior of that cell (not shown). Run 1 for the individually fitted cells has 500 warmup steps and run 2 has 1000. **D**: Comparison of predictions of the dynamics of an unfitted cell using: samples from the posterior distribution of a fitted cell with similar gene expression, samples from the posterior distribution of a random fitted cell, and reference values from literature (“Lemon” values; see Table 1). **E**: Comparison of the effects of HMC parameters. Parameters (*num. warmup steps*; *max. tree depth*) are for r1: (500, 10), r2: (1000, 15), r3: (500, 15).

Parameter inference of the ODE model via a cell chain (denoted *Similar-r1*) was more efficient and gave more accurate results than individually fit cells (Figure 1B–C), with shorter overall computational run times and higher posterior model probabilities (Table S2). The model fit quality was also higher for the cell chain vs. individually fit cells as assessed by the *R* statistic (Table S3). To test whether these improved model fits are in part due to longer fitting times rather than the construction of the cell chain directly, we fit the same cell consecutively ten times: the fits improved over the ten repeat epochs, but the only substantial improvements were seen for the first couple of epochs, after which improvements were minimal and the overall fit quality was comparable to the same cell fitted in the chain (Section S1.2, Figure S4A–F), albeit with some evidence for overfitting in individual parameters (Figure S4G–H). Thus, the quality of fits obtained from fitting in a cell chain are not inherently due to more time spent running inference but are due to the transfer of information between different cells along the chain.

The advantage of transfer learning in cell chains can be demonstrated by the higher predictive power of sampled posterior distributions. We predicted Ca^2+^ responses of test cells for which the parameters have not been inferred (i.e. cells not in *Similar-r1*), using the initial conditions of the test cells but parameters from elsewhere. Each test cell was simulated using parameters sampled from: the posterior of a fitted cell with similar gene expression to the test cell; the posterior of a random fitted cell; and reference values from literature (“Lemon” values; see Table 1). To compare predicted Ca^2+^ responses, we used the Euclidean distance between a predicted Ca^2+^ trajectory and data to quantify the prediction error. We found that posteriors of similar cells and posteriors of random cells had equally good predictive power on test cells: in both cases better than using Lemon values for prediction (Figure 1D). These results illustrate how constructing priors for cells using posterior information from other cells offers greater ability to capture the dynamics of a new cell not previously modeled.

To assess whether cell chains ordered using gene expression information improve inference performance over cell chains ordered randomly, we compared inference runs of at least 500 cells in a chain, with priors informed by cell predecessor, where the chain construction was either random or gene expression similarity-based. The performance of cell chains ordered randomly — evaluated by computational efficiency (sampling times) and accuracy of fits (model posterior probabilities) — was not significantly different than that of the similarity-based chains (Table S4). Therefore, although the use of a cell chain (priors informed by cell predecessors) improved inference relative to individually fit cells, the choice of cell predecessors (similarity-based vs. randomly assigned) did not affect computational efficiency or the accuracy of fits.

We also studied the effects of inference parameters on sampling. We found that sampling times were faster without loss of fit quality when we reduced the maximum tree depth (a parameter controlling the size of the search space) from 15 to 10, since rarely was a tree depth > 10 used in practice; so this reduction did not negatively impact the model fits (Table S5). We also found that a warmup period of 500 steps was sufficient for convergence of MCMC chains for most cells. Setting the maximum tree depth to 10 and the number of warmup steps to 500 led to much faster sampling times for large populations of cells (Figure 1E).

### 3.2 Analysis of single-cell posteriors reveals divergent intracellular and intercellular sources of variability

The posterior distributions of hundreds of cells show striking differences between marginal parameters: some are consistent across cells in a chain while others vary widely. To quantitatively assess this, we ran two cell chains with identical cell ordering for the final 100 cells but with different initial cells. We found that while some marginal posterior parameters were similar for all cells (e.g. *K*_off, ATP_, Figure 2A), others diverged for the same set of cells along a chain (e.g. *d*_5_, Figure 2B). Relative changes in marginal posteriors were seen to be tightly correlated. We computed the fold change in mean marginal posterior parameter values between consecutive cells along the chain (Figure 2A–B, second row): the majority of consecutive cell pairs were tightly correlated both in direction and magnitude, even when the absolute values diverged. We obtained similar results for random cell chains run in parallel with different initial cells (Figure S5). Analysis of the posterior values of parameters relative to their prior distribution revealed that in some cases large deviations were observed. There are several possible causes for large distances existing between prior and posterior distributions, including differences in the biological system (e.g. use of different cell lines) and differences in experimental inputs, e.g. the stimulus used or the amount of stimulus that cells receive.

**Figure 2:**
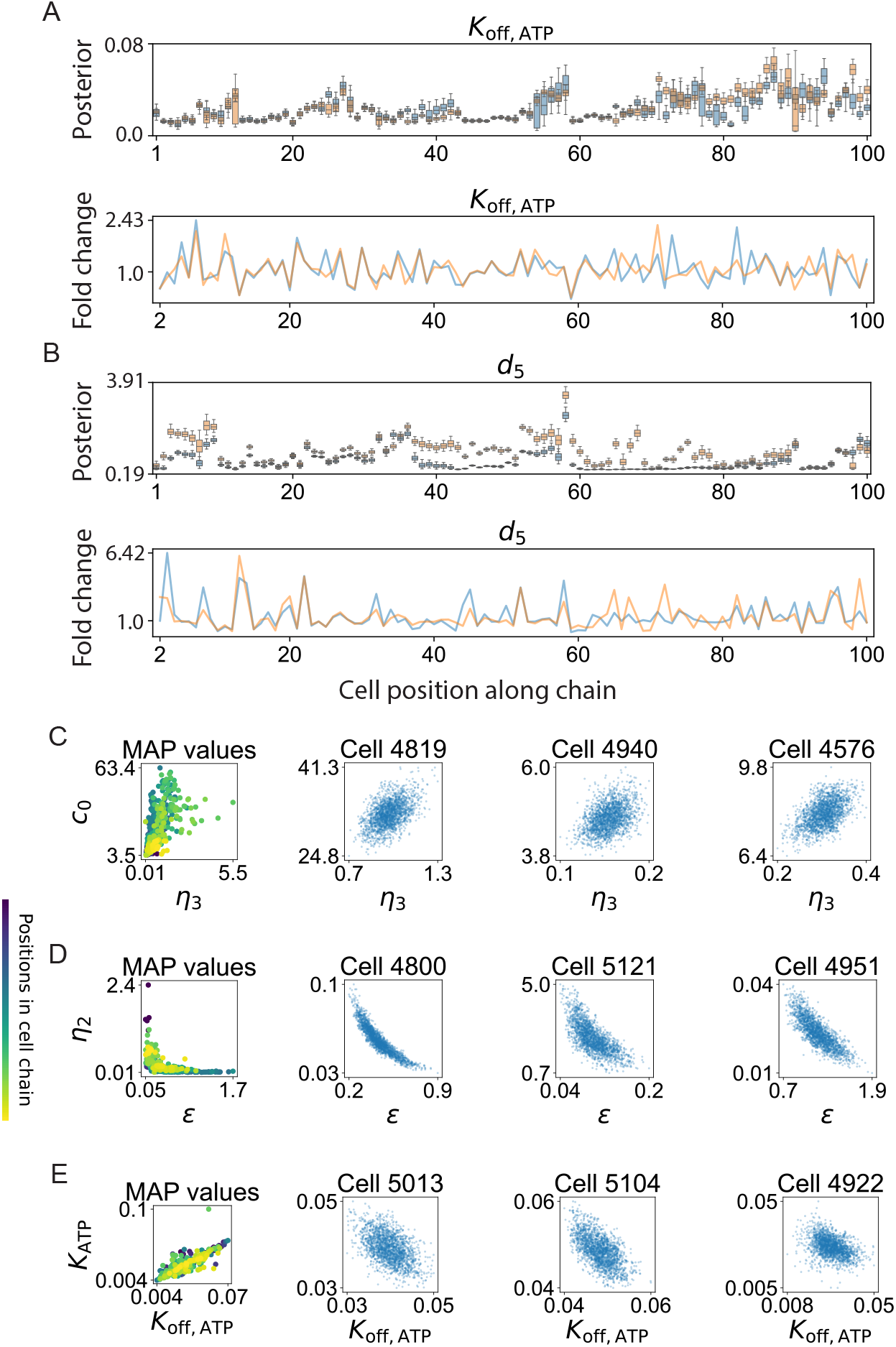
Parameter dependencies revealed by the analysis of marginal posterior distributions along cell chains. **A**: Marginal parameter posterior distributions for the PLC degradation rate (*K*_off, ATP_) in two parallel runs of the same cell chain. Box plots of the first–third quartiles of the distribution with whiskers denoting its full range (upper). Marginal posterior mean fold changes of consecutive cells in the chain (lower). **B**: As for (A), with the IP3 channel dissociation parameter (*d*_5_). **C**: Left: intercellular variability. Scatter plot of the maximum a posteriori (MAP) values of 500 cells for parameters *η*_3_ and *c*_0_. Color indicates position along the chain. Right: intracellular variability. Scatter plots of 500 samples from the posterior of one cell from the chain; three representative cells are shown. **D**: As for (C), with parameters *ϵ* and *η*_2_. **E**: As for (C), with parameters *K*_off, ATP_ and *K*_ATP_.

Further analysis of the marginal posterior distributions revealed two uninformative (“sloppy” [49]) parameters. The posterior distributions of *B*_*e*_ and *η*_1_ drifted, i.e. varied along the chain independent of the particular cell (Figure S6A–B). Given these insensitivities, we studied model variants where either one or both of these parameters were set to a constant. Comparing chains of 500 cells each, the reduced models performed as well as the original in terms of sampling efficiency and convergence (Figure S6C–E, Table S6). Posterior predictive checks of the reduced models showed no significant differences in simulated Ca^2+^ trajectories. Thus, for further investigation into the parameters underlying single-cell Ca^2+^ dynamics, we analyzed the model with both *B*_*e*_ and *η*_1_ set to a constant. This cell chain is referred to as *Reduced-3*.

We discovered striking differences between intracellular and intercellular variability through analysis of the joint posterior distributions of parameters in chain *Reduced-3*. Several parameter pairs were highly correlated, as can be expected given their roles in the Ca^2+^ pathway, e.g. as activators or inhibitors of the same species. However, comparison of parameter correlations within (intra) and between (inter) cells yielded stark differences. Some parameter pairs showed consistent directions of correlation intercellularly (along the chain) and within single cells. The Ca^2+^ pump permeability (*η*_3_) and the concentration of free Ca^2+^ (*c*_0_) were positively correlated both inter- and intracellularly (Figure 2C). Similarly, the ER-to-cytosolic volume (*ϵ*) and the ER permeability (*η*_2_) were negatively correlated in both cases (Figure 2D). However, the ATP decay rate (*K*_ATP_) and the PLC degradation rate (*K*_off, ATP_) were positively correlated along the chain (posterior means) but — for many cells — negatively correlated within the cell (Figure 2E). The distribution of MAP values is well-mixed, i.e. there is no evidence of biases arising due to a cell’s position in the chain: the variation observed in the posterior distributions represents biological differences in the population. These differences may be in part explained by the differences in scale: intercellular parameter ranges are necessarily as large as (and sometimes many times larger than) intracellular ranges. On these different scales, parameters can be positively correlated over the large scale but negatively correlated locally, or vice versa. These divergent sources of variability at the inter- and intracellular levels highlight the complexity of the dynamics arising from a relatively simple model of Ca^2+^ pathway activation.

### 3.3 Quantifying the sensitivity of Ca^2+^ responses in a population of heterogeneous single cells

We conducted analysis of the sensitivity of Ca^2+^ responses to the model parameters. Typically, one defines a parameter sensitivity as the derivative of state variables with respect to that parameter [50, 51]. Here, we are most interested in how the dynamics are affected by parameter perturbations over the range of their marginal posterior distributions. Thus, we evaluate the model response to a given parameter perturbation across its marginal posterior distribution in a population of cells as follows. First, sample from a cell’s posterior distribution, and alter each sample such that the parameter of interest is set to an extreme value according to its marginal posterior distribution (0.01-quantile or 0.99-quantile). We then simulate trajectories from these altered samples (Figure 3A), and use the distance between unperturbed and perturbed model trajectories to define the sensitivity of model output to that parameter, taking the mean of nine simulated trajectories.

**Figure 3:**
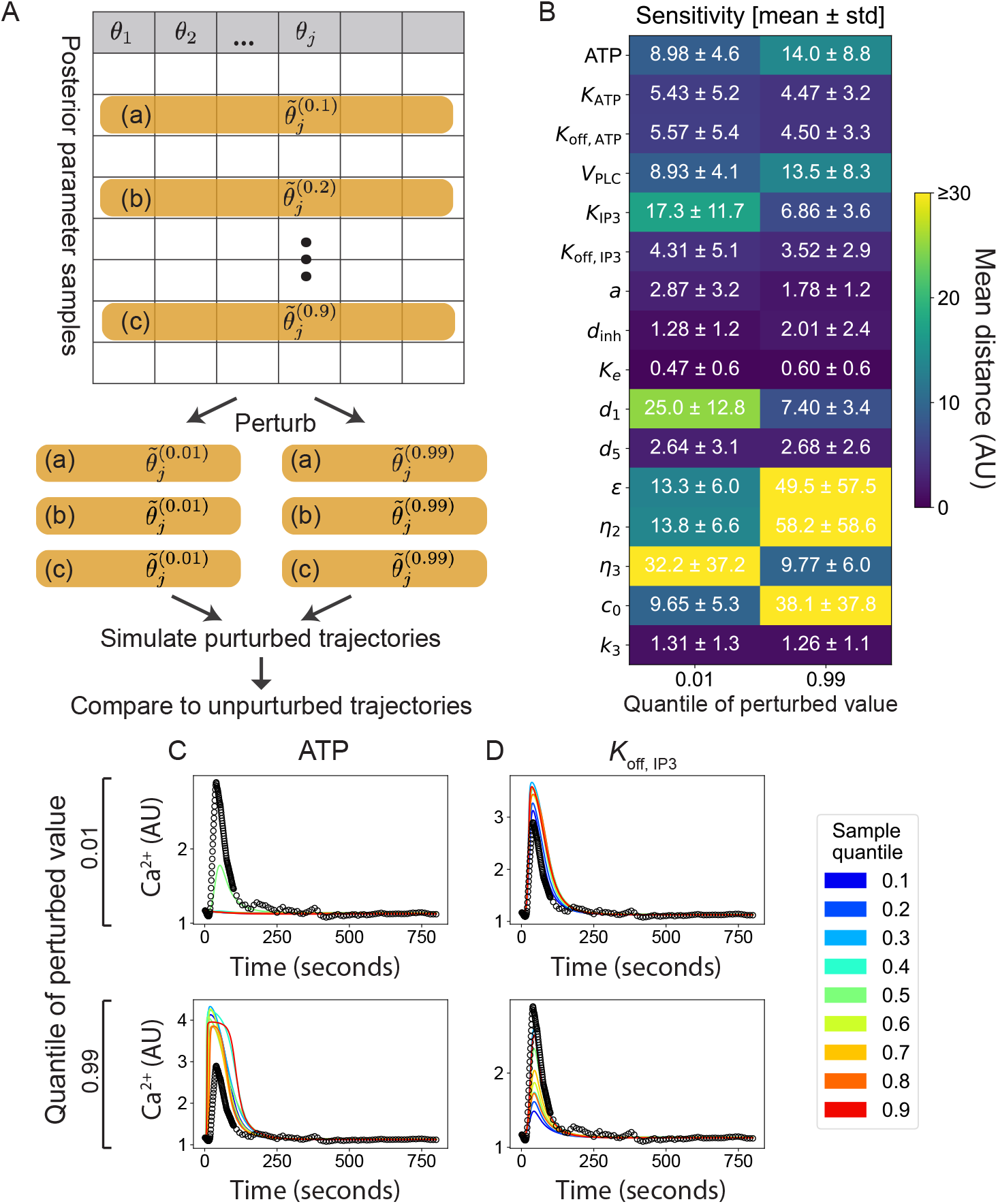
Sensitivity of Ca^2+^ responses to parameter perturbations. **A**: Schematic diagram of approach to studying model sensitivities with respect to parameters sampled across the posterior range. The parameter to be perturbed (*θ*_*j*_) is set to extreme values (0.01- or 0.99-quantile of the marginal distribution). **B**: Parameter sensitivities for a population of fitted cells from a gene expression similarity-based chain (*Reduced-3* model). Sensitivities are reported in terms of model distances between baseline and perturbed trajectories. std: standard deviation. **C**: Simulated model trajectories in response to perturbations in ATP concentration. **D**: Simulated model trajectories in response to perturbations in *K*_off, IP3_.

We find that there is a lot of variation in the Ca^2+^ responses: sensitive to some model parameters and insensitive to others (Figure 3B). Notably, the sensitivities of the least sensitive parameters had mean values of close to 1.0: similar to the distances obtained from the best-fit posterior values (Table S5), i.e. the Ca^2+^ response is insensitive to these parameters across the whole posterior range. The insensitive parameters were not simply those which had the highest posterior variance: there was little correlation between the inferred sensitivity and the posterior variance (Table S7), compare, e.g., parameters *d*_1_ and *d*_5_.

Analysis of the Ca^2+^ responses to parameter perturbations provides means to predict how much Ca^2+^ responses are affected by changes in extracellular and intracellular dynamics (Figure 3C–D). For example, low concentrations of ATP result in very low Ca^2+^ responses; increasing the concentration of ATP can more than double the peak response (Figure 3C). The importance of IP3 in Ca^2+^ signal transduction is in agreement with the results of Yao et al. [38]; here we go further in that we can quantify the particular properties of the Ca^2+^ response affected by each parameter. In the case of *K*_off, IP3_, the main effect is also in the peak height of the Ca^2+^ response (Figure 3D).

### 3.4 Variability in gene expression is associated with variability in Ca^2+^ dynamics

We studied variation between pairs of genes and parameters sampled from a cell population to assess whether relationships between them might exist. We found that several gene-parameter pairs were correlated. In general, the proportion of variance explained between a gene-parameter pair was low; this is to be expected given the many sources of variability in both the single-cell gene expression and the Ca^2+^ responses.

Analysis of the most highly correlated gene-parameter pairs (see Methods and Table S8) identified a number of genes that were correlated with multiple parameters, e.g. PPP1CC, as well as parameters that were correlated with multiple genes, e.g. *η*_3_. Pairwise relationships were analyzed via linear regression. The top four correlated gene-parameter pairs from a similarity-based cell chain are shown in Figure 4A–D: cells are well-mixed according to their positions along the chain, i.e. correlations are not due to local effects. The pairwise correlations overall are low, which we expect given single-gene inputs. Performing multiple regression could improve predictive power, however our goal here is to study whether any evidence supports the existence of individual gene-parameter relationships. We performed the same analysis on a randomly ordered cell chain, where the same gene-parameter relationships were recapitulated, albeit with lower absolute correlation values (Figure 4E–H and Table S9). There is no discernable influence of a cell’s position in a chain on the gene-parameter relationship, confirming that these correlations among a cell population reflects the variability in the population rather than any sampling artefacts.

**Figure 4:**
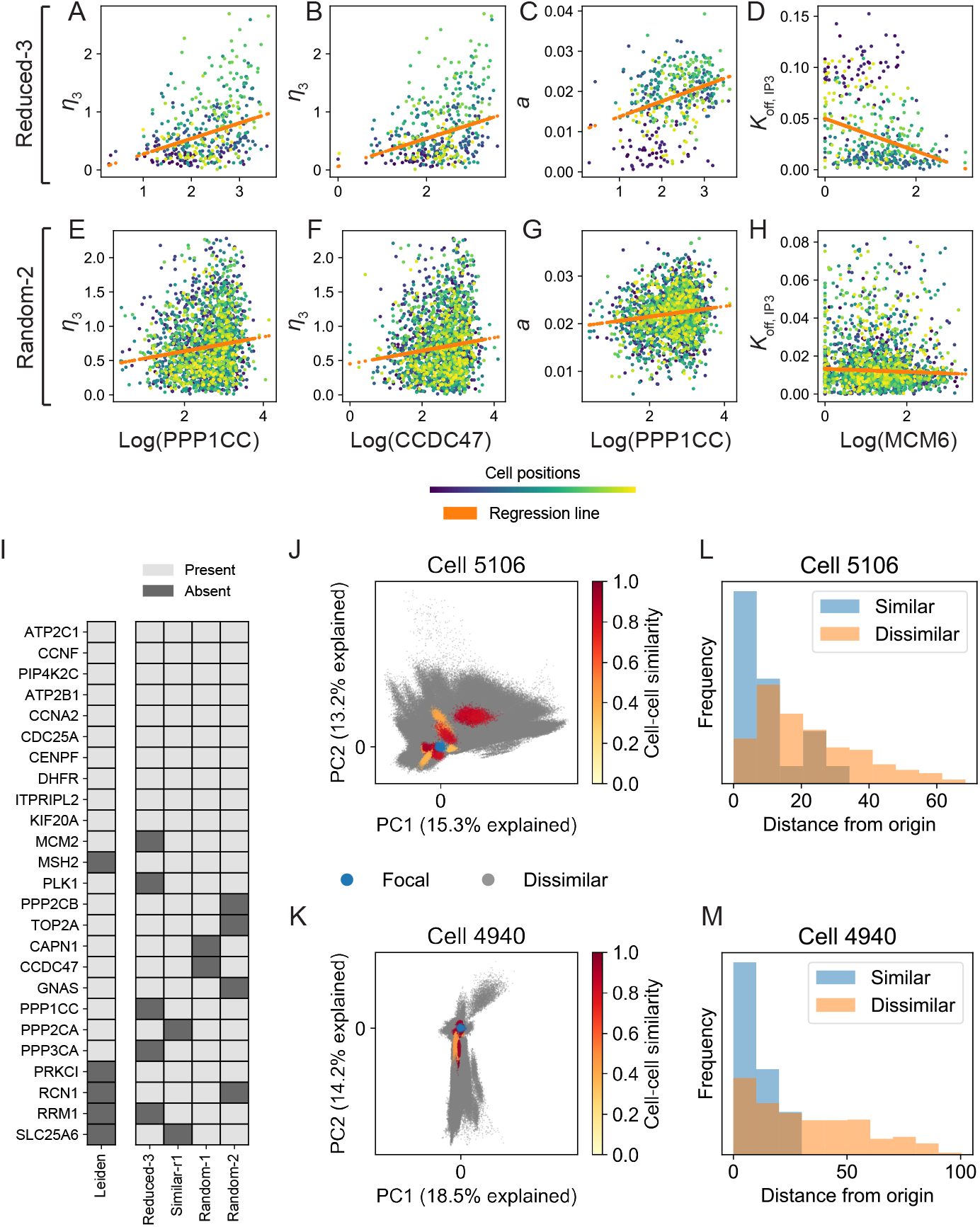
Variability in Ca^2+^ model dynamics is associated with variability in gene expression. **A–D**: Top ranked gene-parameter pairs by Pearson correlation from similarity-based chain (*Reduced-3*). Log gene expression is plotted against marginal posterior means. Each dot represents a cell; color denotes cell position along the chain. **E–H**: The same gene-parameter pairs shown in (A), for cells sampled from a randomly ordered cell chain (*Random-2*). **I**: Comparison of genes identified as present/absent in the top 30 genes from gene-parameter correlation analysis across different chains, and the top 30 marker genes from Leiden clustering directly on the gene expression space. **J**: Projection of posterior samples (500 per cell) onto the first two principal components of the focal cell (Cell 5106), shown in blue. Posterior samples from dissimilar cells shown in gray; posterior samples from similar cells shown in yellow-red, where the shade indicates degree of transcriptional similarity. **K**: As for (J), with PCA performed on a different focal cell (Cell 4940). **L**: Histogram corresponding to (J): mean distances from the origin (focal cell 5106) to samples from surrounding cells. **M**: Histogram corresponding to (K): mean distances from the origin (focal cell 4940) to samples from surrounding cells.

We compared the top genes ranked by gene-parameter correlations for four populations: from two randomly sampled and two similarity-informed cell chains. Gene-parameter pairs were sorted by their absolute Pearson correlation coefficients, and the genes ranked by their positions among sorted pairs. In total we identified 75 correlated gene-parameter pairs for the *Reduced-3* chain, applying a Bonferroni correction for multiple testing (Figure S7). Out of the top 30 of these, 25 appeared in the top 30 in at least 3/4 of the cell chains studied (Figure 4I). Of these 25 genes, 20 also appeared as top-10 marker genes from unsupervised clustering (into 3 clusters) of the gene expression data directly (Figure 4I). The high degree of overlap between these gene sets demonstrates that a subset of genes expressed in MCF10A cells explain not only their overall transcriptional variability but also their variability in Ca^2+^ model dynamics. These results are also suggestive of how information content pertaining to the heterogeneous Ca^2+^ cellular responses is encapsulated in the parameter posterior distributions.

Next, we turn our attention from the level of individual genes/parameters to that of the whole: what is the relationship between the posterior parameter distribution of a cell and its global transcriptional state? We used principal component analysis (PCA) for dimensionality reduction of the posterior distributions to address this question. We selected a cell (denoted the “focal cell”) from a similarity-based cell chain (*Reduced-3*) and decomposed its posterior distribution using PCA. We projected the posterior distributions of other cells onto the first two components of the focal cell (Figure 4J–K and Figure S8A–B) to evaluate the overall similarity between the posterior distributions of cells relative to the focal cell. On PCA projection plots, posterior samples are colored based on gene expression: samples are derived from cells that are either transcriptionally similar to the focal cell, or share no transcriptional similarity. Comparison of similar and dissimilar cells from the same population showed that cells that were transcriptionally similar were located closer to the focal cell than dissimilar cells (Figure 4L–M and Figure S8C–D). In contrast, similar analysis of a random cell chain showed that transcriptionally similar cells were not located closer to the focal cell than dissimilar cells (Figure S9). Notably, proximity of posterior samples derived from transcriptionally similar cells was not driven by a cell’s position along the chain (no block structure observed; Figure S10). Similarities between posterior distributions of transcriptionally similar cells were thus not driven by local cell-cell similarity, but rather underlie a global effect and denote a relationship between the transcriptional states of cells and the Ca^2+^ pathway dynamics that they produce.

### 3.5 Similarity-based posterior cell clustering reveals distinct transcriptional states underlying Ca^2+^ dynamics

To characterize the extent to which we can predict Ca^2+^ responses from knowledge of the model dynamics, we clustered 500 cells from a similarity-based cell chain (*Reduced-3*) based on the single-cell posterior distributions using hierarchical clustering (see Methods). Three clusters were obtained (Figure 5A). Each cluster showed distinct Ca^2+^ dynamics: “low-responders” exhibited lower overall Ca^2+^ peaks in response to ATP (Figure 5B); “early-responders” exhibited earlier overall Ca^2+^ peaks in response to ATP; and “late-high-responders” exhibited robust Ca^2+^ responses with peaks that were later and higher than cells from other clusters (Figure S11). The distinct dynamic profiles can be explained by the model parameters that give rise to them: low-responders were characterized by high concentration of free Ca^2+^ in the ER (*c*_0_) and low activation rates of IP_3_R (Figure S11, Figure S12, Figure S14). Early-responders were characterized by parameters leading to faster and earlier IP3 and PLC dynamics. Late-high-responders were characterized by small values of *d*_1_ (Figure S14).

**Figure 5:**
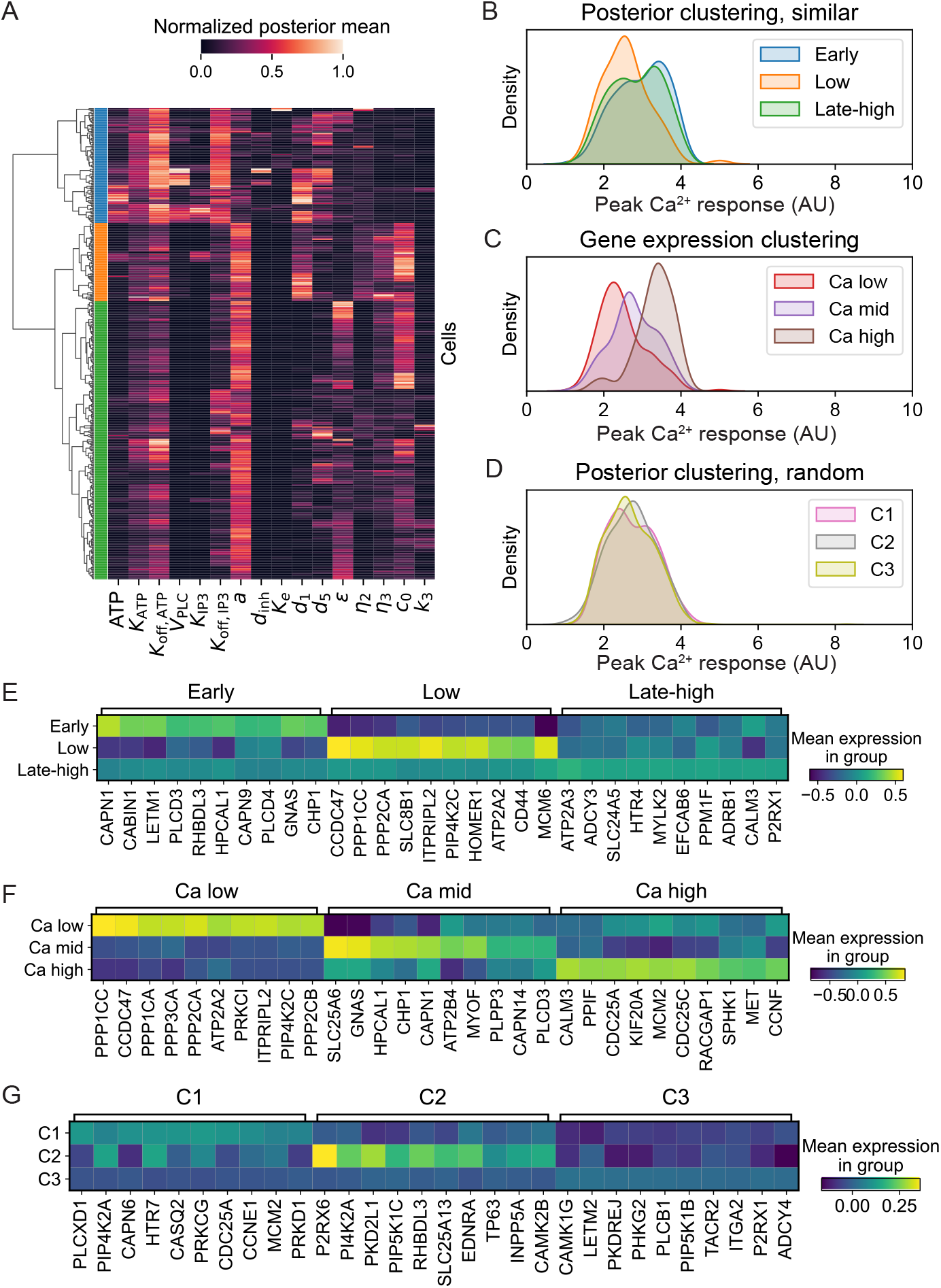
Clustering of cell posterior distributions reveals marker genes for Ca^2+^ states. **A**: Agglomerative clustering on posterior means from a similarity-based chain (*Reduced-3*) using Ward linkage (*k* = 3 clusters). **B**: Kernel density estimate of Ca^2+^ peak height from posterior clustering of a similarity-based chain. **C**: Kernel density estimate of Ca^2+^ peak height from gene expression clustering (Leiden) for the same cell population as shown in (A–B). **D**: Kernel density estimate of Ca^2+^ peak height from posterior clustering of a randomly ordered cell chain (*Random-2*). **E**: Top ten marker genes per cluster from similarity-based posterior clustering. **F**: Top ten marker genes per cluster from gene expression clustering of cells. **G**: Top ten marker genes per cluster from posterior clustering of randomly ordered cell chain.

Comparison of posterior parameter clustering with the clustering done by Yao et al. [38] highlights similarities and distinctions. In both cases, one of the three clusters was characterized by larger responses to ATP and correspondingly higher values of *d*_inh_ (Figure S14). In Yao et al., both *d*_1_ and *d*_5_ were smaller in cells with stronger Ca^2+^ responses; we found that *d*_1_ was smaller in the late-high-responder cluster, but not in the early responders. In our results, *d*_5_ was higher for the early-responders, in contrast with Yao et al. (Figure S14). We note that we set a stringent threshold for minimum peak Ca^2+^ response, i.e. we excluded non-responding cells, unlike Yao et al., thus in a direct comparison most of the cells in our population would belong to the “strong positive” cluster in [38].

To assess the Ca^2+^ dynamic clusters we obtain in light of single-cell gene expression, we performed two analyses for comparison. We clustered the same 500 cells based solely on their gene expression using community detection (Leiden algorithm in Scanpy [46]); and we clustered cells from a randomly ordered cell chain using the same approach as above for hierarchical clustering of posterior parameters. The cell clusters obtained based solely on gene expression can be distinguished based on the Ca^2+^ profiles observed: “Ca-low”, “Ca-mid”, and “Ca-high” responses (Figure 5C); overlapping partially with the similarity-based clusters obtained (Figure 5B). In contrast, no distinct Ca^2+^ dynamic responses could be observed for the posterior clustering based on the random cell chain (Figure 5D and Figure S13).

We analyzed differential gene expression from each set of clusters obtained, from the similarity-based chain (Figure 5E), the gene expression-based clustering (Figure 5F), and the randomly ordered chain (Figure 5G). Distinct markers for each cluster were observed for the similarity-based clustering and the gene expression-based clustering, but were not discernible for cell clustering on the random chain. Clustering of cell posteriors from the randomly ordered chain was thus unable to distinguishable Ca^2+^ dynamic profiles nor gene expression differences. On the other hand, clustering posteriors from a similarity-based chain identified distinct gene expression profiles, and these overlapped with the marker gene profiles obtained by clustering on the gene expression directly. I.e. parameter inference of single-cell Ca^2+^ dynamics from using a gene expression similarity-based chain enables the identification of cell clusters with distinct transcriptional profiles and distinct responses to ATP stimulation.

Analysis of the genes that are associated with each Ca^2+^ profile showed that low-responder cells were characterized by upregulation of CCDC47 and PP1 family genes (PPP1CC and PPP2CA). Early-responder cells were characterized by upregulation of CAPN1 and CHP1, among others. The late-high responder cells were characterized by increased expression CALM3 among others, although the marker genes for this cluster were less evident than the others. By comparison of marker genes, we saw considerable overlap between the early-responders (similarity-based clustering) and the Ca-mid responders (gene expression clustering). We also saw overlap between the low-responder and the Ca-low marker gene signatures. These results highlight that the posterior distributions of cells fit from similarity-based cell chains captured information about the underlying transcriptional states of the cells, linking cellular response parameters directly to transcriptional states. For example, the low-responder cells (similarity-based clustering) were distinguished by parameter *d*_5_, characterizing the dynamics of IP3 dissociation, and were marked by high PPP1CC and CCDC47 expression.

Finally, we considered whether alternative means for cell chain construction could provide similar information. We constructed a cell chain using cells from *Reduced-3*, denoted “*Ca-similarity*” for which consecutive cells displayed similar Ca^2+^ responses (see Section S1.3). Clustering cells from *Ca-similarity* based on their Ca^2+^ responses (via *k*-means) showed that cells with different Ca^2+^ responses had distinct gene expression profiles (Figure S15). However, hierarchical clustering on the parameter posterior distributions from these cells after performing inference on *Ca-similarity* did not separate cells into clusters with distinct Ca^2+^ responses or distinct gene expression profiles (Figure S16A–B). This result makes sense when analyzed via the similarity matrices obtained for *Ca-similarity* vs. a gene expression similarity-based chain (Figure S16C–D). For the former, almost all pairs of neighboring cells did not share similarity in gene expression despite their similarity in Ca^2+^ response. Whereas for *Reduced-3*, the gene expression similarity-based chain, all pairs of consecutive cells were similar in gene expression (Figure S16D).

## 4 Discussion

We have presented methods for inferring the parameters of a signaling pathway model, given data describing dynamics in single cells coupled with subsequent gene expression profiling. We hypothesized that via transfer learning we could use posterior information from a cell to inform the prior distribution of its neighbor along a “cell chain” of transcriptionally similar cells. Implemented using Hamiltonian Monte Carlo sampling [42], we discovered that the cell chain construction for prior distributions did indeed lead to faster sampling of parameters. However, these improvements did not rely on the use of gene expression to construct priors: the performance of inference on cells in a chain ordered randomly was equally good. However, cell chains constructed using gene expression similarity contained more information (as defined by their posterior parameter distributions) about Ca^2+^ signaling responses. Clustering the posterior parameters identified important relationships between gene expression and the dynamic Ca^2+^ phenotypes, thus providing mappings from state to dynamic cell fate.

The model studied here is described by ODEs to characterize the Ca^2+^ signaling pathway, adapted from [40, 37], consisting of 12 variables and (originally) a 40-dimensional parameter space. This was reduced to 19 parameters in Yao et al. [38] and 16 parameters in our work. Analysis of even a single 16-dimensional posterior distribution requires dimensionality reduction techniques, let alone the analysis of the posterior distributions obtained for populations of hundreds of single cells. Parameter sensitivity analysis highlighted the effects of specific parameter perturbations on the Ca^2+^ dynamic responses. Indeed, we advocate for the use of sensitivity analysis more generally as means to distinguish and pinpoint the effects of different parameter combinations for models of complex biochemical signaling pathways.

By unsupervised clustering of the posterior distributions, we found that distinct patterns of Ca^2+^ in response to ATP could be mapped to specific variation in the single-cell gene expression. In previous work using similar approaches for clustering [38], posterior parameter clusters predominantly revealed response patterns consisting of responders and non-responders; here we excluded those cells that did not exhibit a robust response to ATP. We were able to characterize subtler the Ca^2+^ response dynamics (described by “early”, “low”, and “late-high” responders) and predict which transcriptional states give rise to each. This approach is limited since relatively little gene expression variance is explained by an individual model parameter: it may be possible to address this in future work by surveying a larger range of cell behaviors, e.g. by including a wider range of cellular responses or by considering higher-level co-variance in the posterior parameter space. It also remains to be tested whether the given model of Ca^2+^ dynamics is appropriate to describe the signaling responses in cell types other than MCF10A cells.

Our ability to fit to of the single cells tested came potentially at the expense of an unwieldy model size. With four variables and a 16-dimensional parameter space, the dimension of the model far exceeds that of the data: time series of Ca^2+^ responses in single cells. Without data with which to constrain the three additional model species, we needed to constrain the model in other way. We used an approach of “scaling and clipping” for construction of the priors, i.e. setting ad hoc limits to control posterior variance. More effective (and less ad hoc) techniques could improve inference overall and may become necessary in the case of larger models. These include (in order of sophistication): tailoring the scaling/clipping choices to be parameter-specific; tailoring the choice of prior variance based on additional sources of data; or performing model reduction/identifiability analysis to further constrain the prior space before inference. Constructing priors from cells with similar gene expression also helped to curb the curse of dimensionality: sampling cells sequentially places a constraint on the model. Nonetheless, in the future more directed approaches to tackle model identifiability ought to be considered.

Connecting dynamic cell phenotypes to transcriptional states remains a grand challenge in systems biology. The limitations of deriving knowledge from gene expression data alone [24] have led to the proposal of new methods seeking to bridge the gap between states and fates [52]. Here, making use of technology that jointly measures Ca^2+^ dynamics and gene expression in single cells, we have shown that parameter inference informed by transcriptional similarity represents another way by which we can connect gene expression states to dynamic cellular phenotypes. The statistical framework employed improved our ability to perform parameter inference for single-cell models. We expect that improvements to statistical inference frameworks could be gained by similar approaches applied to other models of nonlinear cellular response dynamics. Current and future technologies combining higher-resolution dynamic cell measurements with high-throughput genomics will provide new data to inform these parameter inference methods and, we expect, lead to new discoveries of means of transcriptional control of biological dynamics.

## Acknowledgments

This work was supported by an Andrew J. Viterbi Fellowship in Computational Biology and Bioinformatics (to X.W.), A.L.M. acknowledges support from the National Institutes of Health (R35GM143019) and the National Science Foundation (DMS2045327).

## Data Availability

Parameter inference was developed in Python 3.6 and Stan 2.19. Posterior analyses were developed in Python 3.8. All code developed to simulate models and run parameter inference is released under an MIT license at: https://github.com/maclean-lab/singlecell-parinf. Ca^2+^ data and MERFISH data (processed files) are also available at the GitHub repository.

## Notes

https://github.com/maclean-lab/singlecell-parinf

### Competing Interest Statement

RW is a co-founder and equity holder of a BioCartography inc. The other authors have no competing interests to declare.

### Summary of Updates

Edits made to figures for clarity and edits to text made throughout to improve readability and discuss new analyses.

