## Supplementary Figures for "Single-cell Ca^2+^ parameter inference reveals how transcriptional states inform dynamic cell responses"

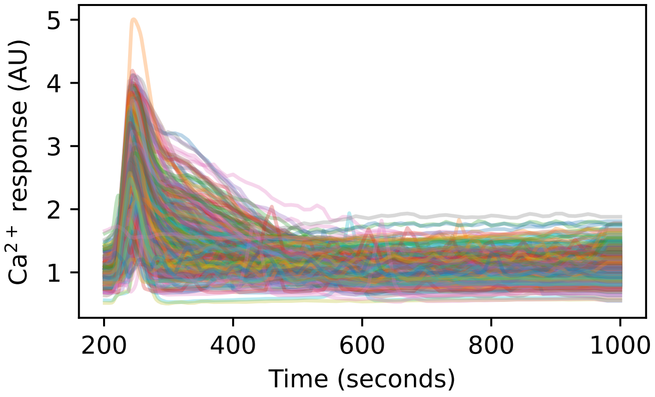


Figure S1: **$Ca^{2+}$ dynamic response trajectories to stimulus by ATP.** Responses for 500 cells are shown over a time frame of 800 seconds. Cells were stimulated by ATP at 200 seconds; the first 200 seconds prior to stimulation are not shown. The raw $Ca^{2+}$ trajectories were smoothed using a moving average filter with a window size of twenty seconds.

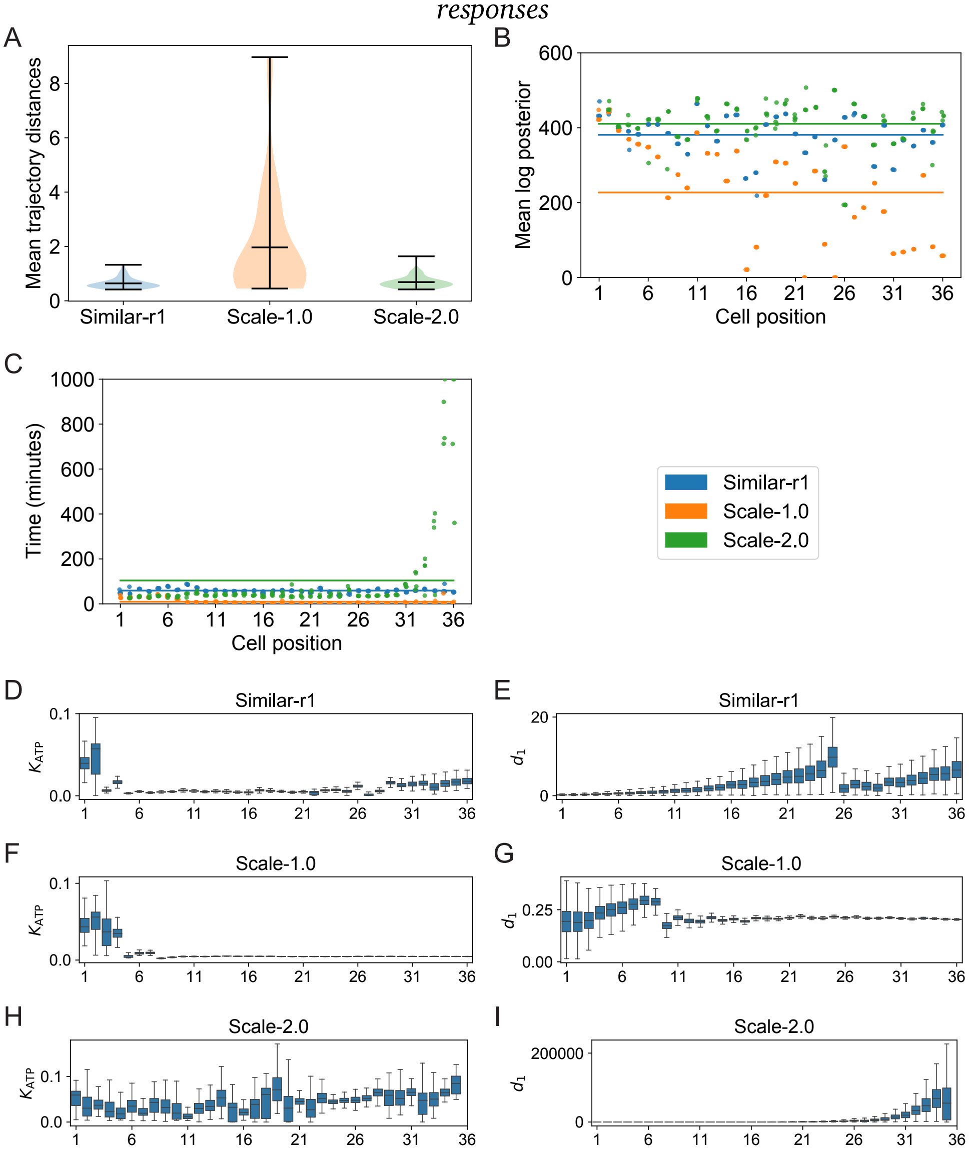


Figure S2: **Analysis of scaling and clipping of prior distributions along cell chains. A**: Mean error of trajectories simulated from parameter posterior of 36 cells sampled from one of three chains, (*Similar-r1*): posterior parameter standard deviation is scaled by a factor of 1.5 and clipped; (*Scale-1.0*): posterior parameter standard deviation scaled by a factor of 1.0; and (*Scale-2.0*): posterior parameter standard deviation scaled by a factor of 2.0. Means and ranges are shown on the violin plots. Only 36 cells are compared per chain because at this point in the chain run times for *Scale-2.0* became prohibitive (see C). **B**: Mean log posterior values per cell for each of the three chains between sampled trajectories and the data. Mean value over a whole chain is given as horizontal line. **C**: Mean sampling times per cell. Mean value over a whole chain is given as horizontal line. **D–I**: Marginal parameter distributions for $K_{\mathrm{ATP}}$ and $d_1$ for *Similar-r1* (D–E), *Scale-1.0* (F-G) and *Scale-2.0* (H–I).

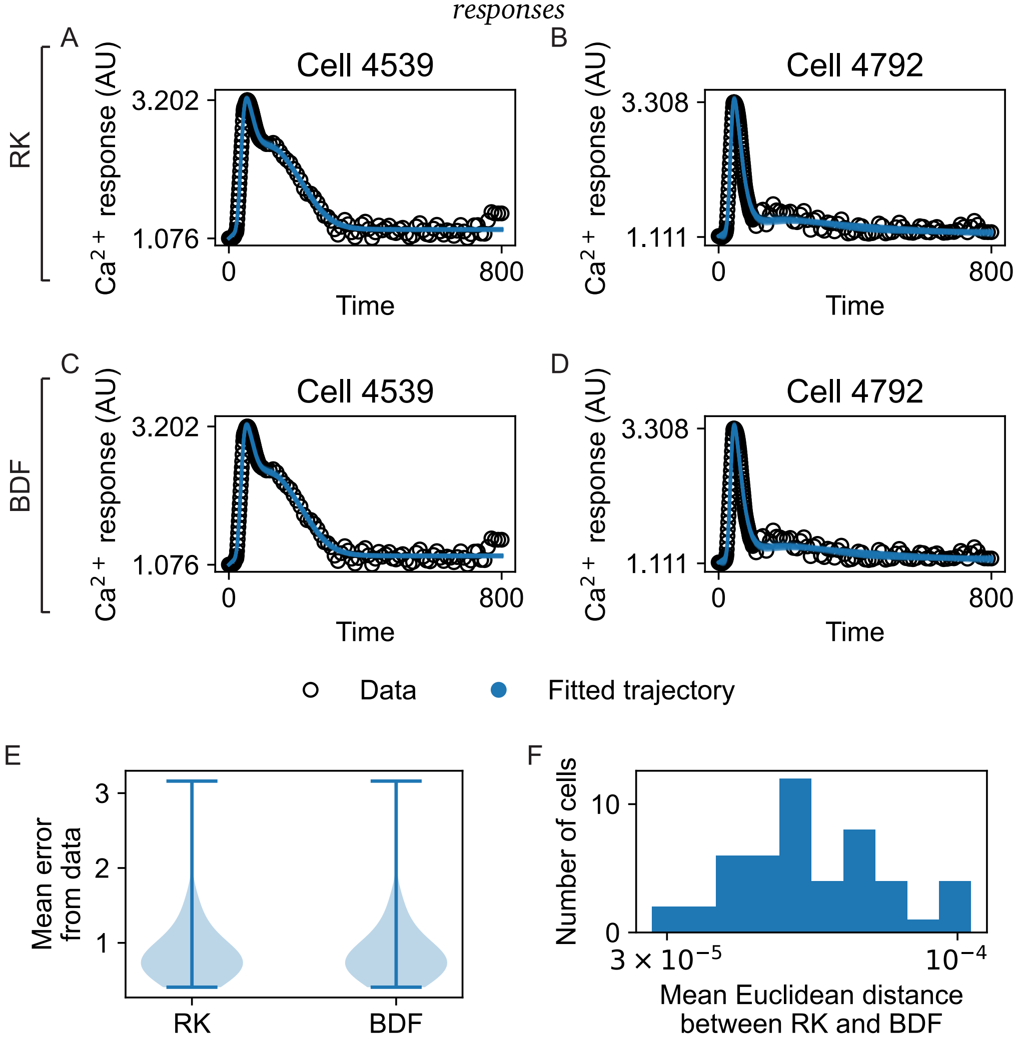


Figure S3: **Comparison of stiff and non-stiff solvers for $Ca^{2+}$ response simulation.** We simulated $Ca^{2+}$ responses from sampled posterior for each individual cell with a non-stiff ODE solver (Runge-Kutta, RK) and a stiff solver (based on backward differentiation formula, BDF). **A–B**: Examples of $Ca^{2+}$ response simulated by a non-stiff solver (RK scheme). **C–D**: Examples of $Ca^{2+}$ response simulated by a stiff solver (BDF scheme). **E**: Mean error from data using RK and BDF solvers for all cells. **F**: Mean Euclidean distance between simulated $Ca^{2+}$ trajectories from both solvers for the same cells.

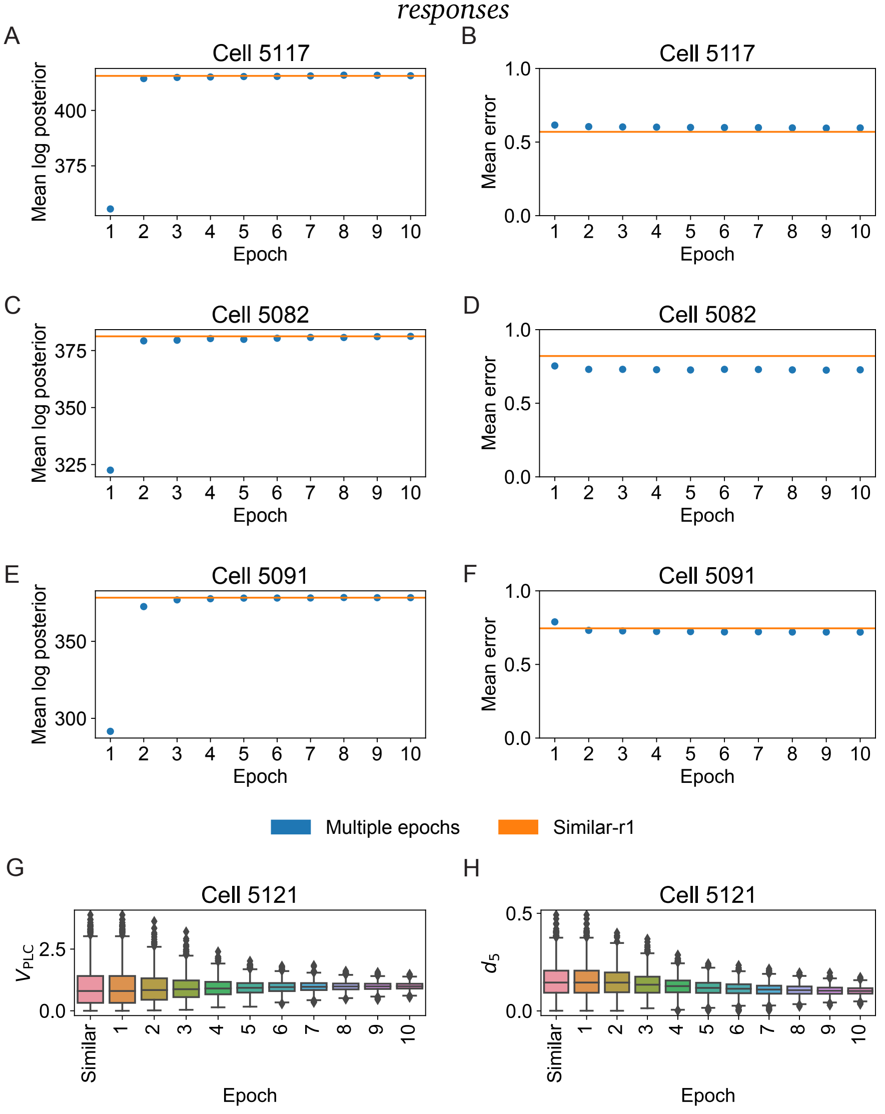


Figure S4: **Comparison of parameter inference of cells in a chain vs. multiple inference runs on the same cell.** We trained individual cells with multiple epochs, starting with the Lemon prior for the first epoch. **A**: Posterior probabilities of sampled parameters for Cell 5117, multi-epoch learning (blue points) vs. the cell as fit in the chain *Similar-r1*. **B**: Mean error of trajectories simulated from sampled posterior parameters for Cell 5117, multi-epoch learning vs. the cell as fit in the chain *Similar-r1*. **C**: As for (A) with Cell 5082. **D**: As for (B) with Cell 5082. **E**: As for (A) with Cell 5091. **F**: As for (B) with Cell 5091. **G**: Marginal posterior distributions of $V_{\text{PLC}}$ from multi-epoch training for cell 5121 vs. the cell as fit in the chain *Similar-r1* (“Similar”). **H**: As for (G) with $d_5$.

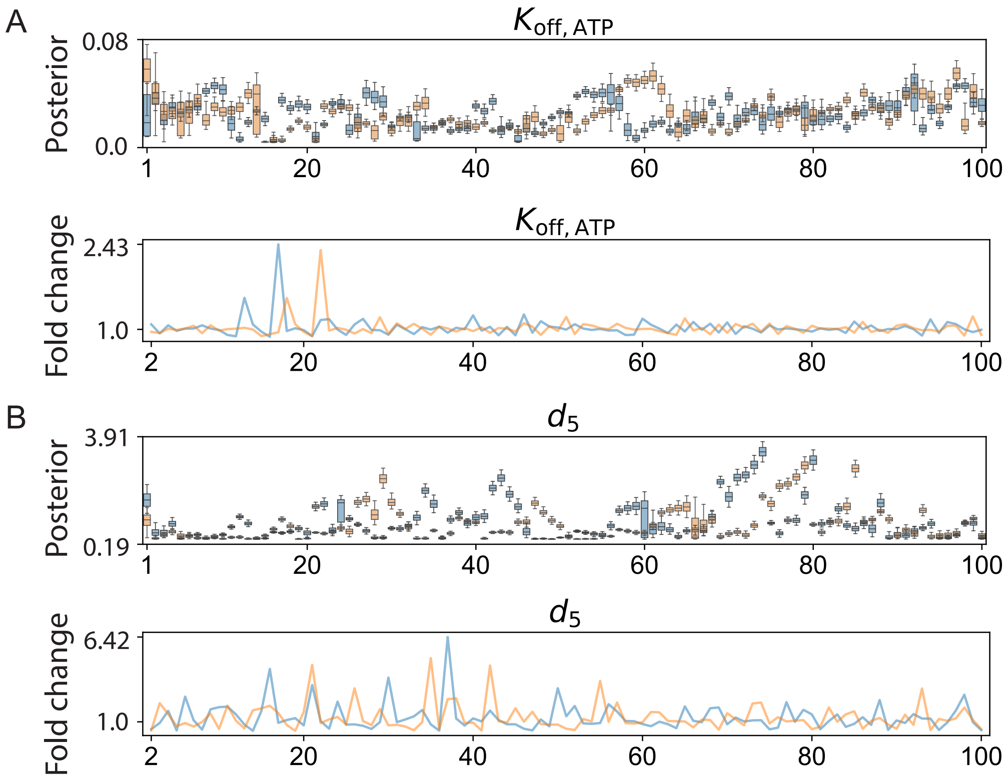


Figure S5: **Comparison of replicate runs of random cell chains. A**: Two randomly ordered cell chains were run with different initial cells (6 unique initial cells for the blue chain and 3 for the orange) followed by the same 100 cells. Marginal parameter posterior distributions for $K_{\text{off, ATP}}$ (PLC degradation rate) are shown (top), and the corresponding fold changes between consecutive cells (bottom). **B**: Marginal posteriors and fold changes as for (A), with $d_5$, the IP3 channel dissociation constant.

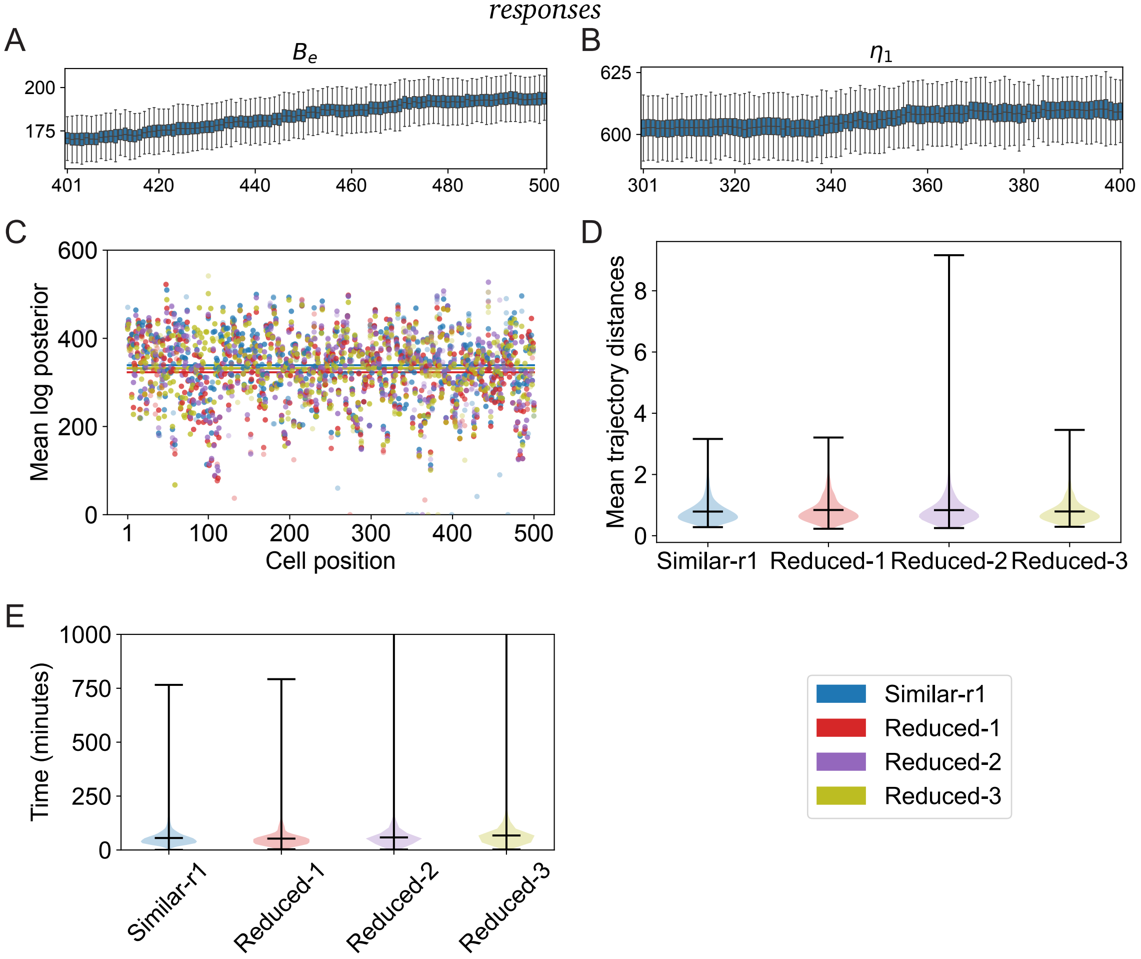


Figure S6: **Comparison of model reduction for insensitive parameters. A–B**: Marginal posterior values for $B_e$ (A) and for $\eta_1$ (B) from the chain *Similar-r1*. In both cases 'drift' is observed over 100 fitted cells (positions 401–500 in the cell chain in (A) and 301–400 for (B)): these parameters are insensitive given the data. **C**: Comparison of the full model (*Similar-r1*) with three reduced models. In *Reduced-1*, $\eta_1$ is set to a constant value; in *Reduced-2*, $B_e$ is set to a constant value; in *Reduced-3*, both $\eta_1$ and $B_e$ are set to constant values. Constants taken from previous parameter estimation in Lemon et al. [37]. Mean log posterior values are shown for all 500 cells. Means over cell chain is shown as horizontal line. No significant differences are observed. **D**: Mean errors between sampled trajectories and the data. **E**: Mean sampling times for each of the four chains. No significant differences are observed.

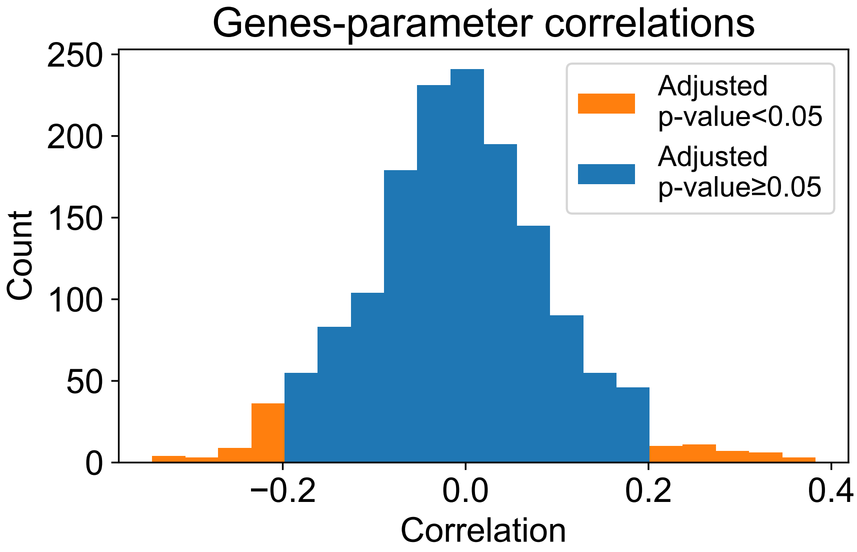


Figure S7: **Distribution of Pearson correlations between genes and parameters from the chain *Reduced-3*.** For all significant gene-parameter pairs (adjusted p-value $< 0.05$), the absolute Pearson correlation was $> 0.2$.

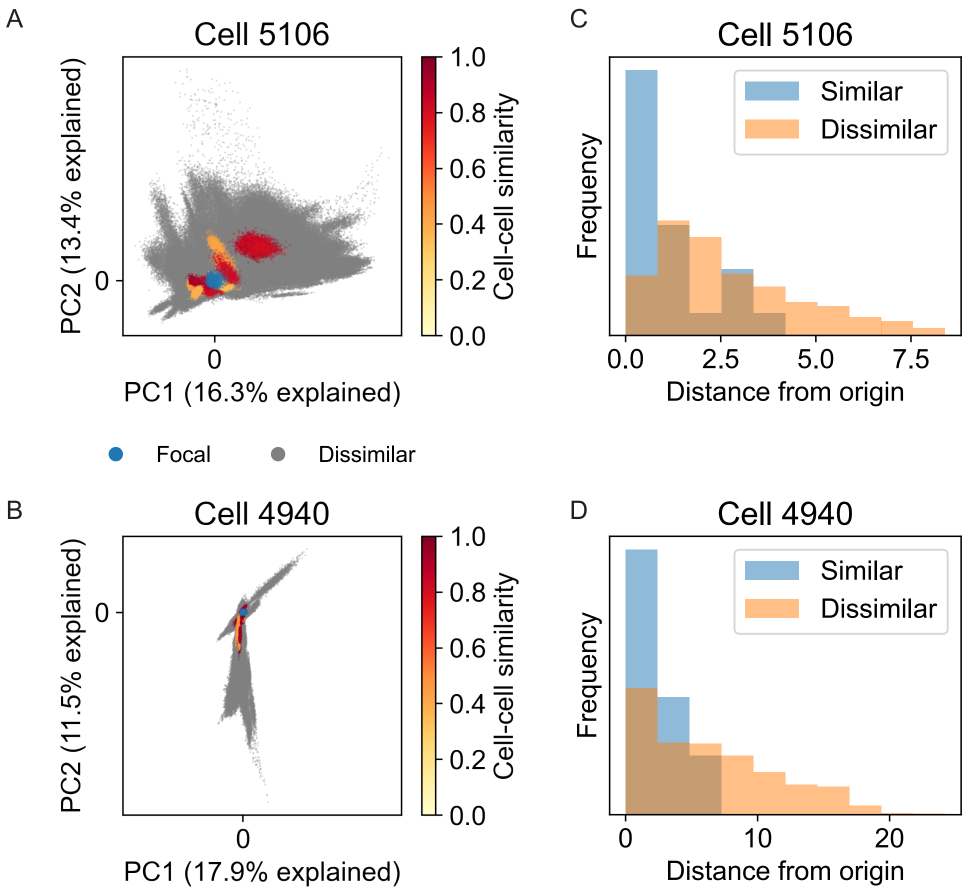


Figure S8: **Comparison of global similarity via PCA using min-max normalization for chain *Reduced-3*. A**: Projection of all cells from the *Reduced-3* chain onto first two PCs of cell 5106 via min-max normalization (cf. z-score normalization used in Fig 4). **B**: As for (A), with focal cell 4940. **C**: Mean distances between focal cell 5106 and projected samples from (A). **D**: Mean distances between focal cell 4940 and projected samples from (B).

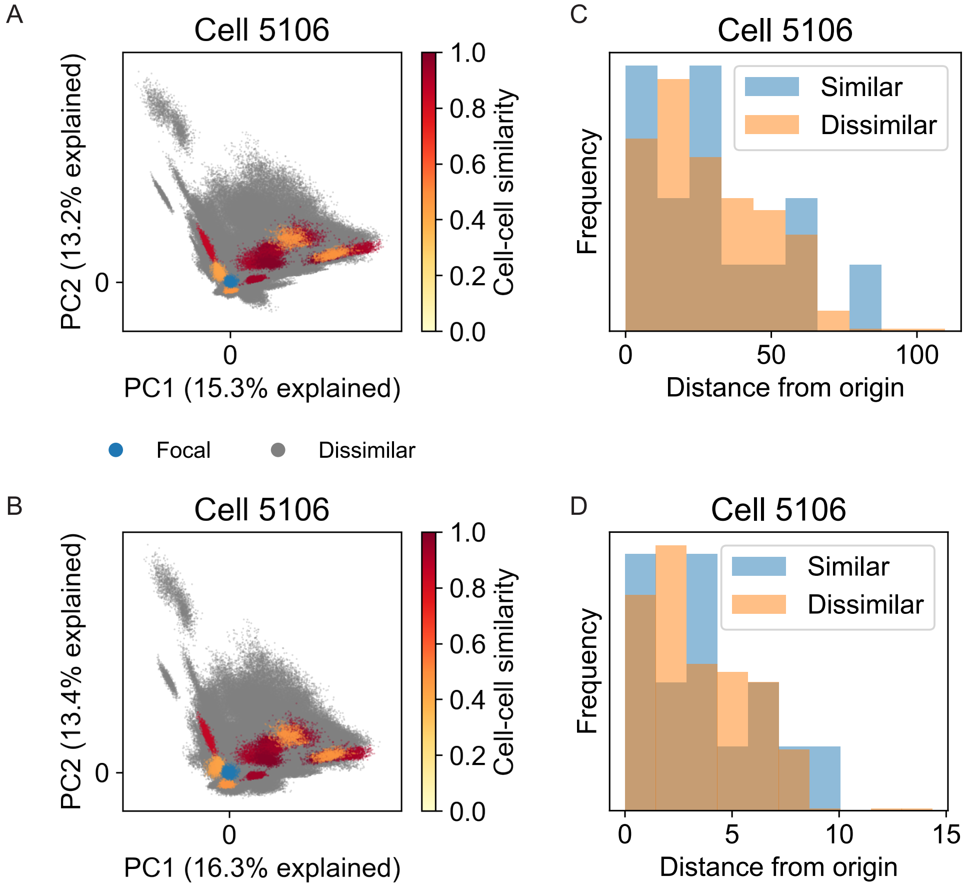


Figure S9: **Comparison of global similarity via PCA using min-max normalization for chain *Random-2*.** **A**: Projection of all cells from the *Random-2* chain onto first two PCs of cell 5106 under $z$-score normalization. **B**: Projection of all cells onto first two PCs of cell 5106 under min-max normalization. **C**: Mean distances between focal cell 5106 and projected samples from (A). **D**: Mean distances between focal cell 5106 and projected samples from (B).

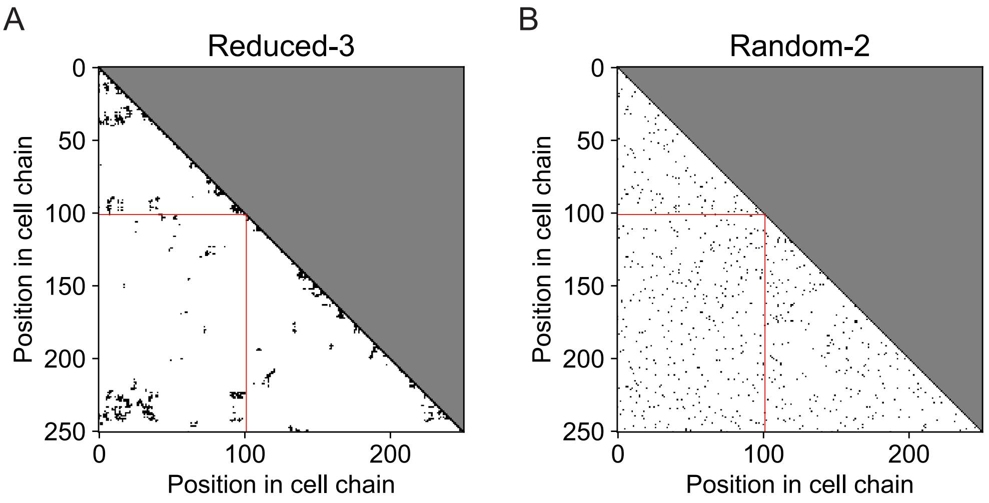


Figure S10: **Comparison of cell-cell similarity along cell chains. A**: Black spots in the lower triangle indicate that the cell pair has gene expression similarity $> 0$ for the *Reduced-3* chain. Red lines mark the row/column of the $101^{st}$ cell. Small clusters of similar cells are observed however these do not locate preferentially near the diagonal, i.e. we do not see evidence of block structure. **B**: As for (A), for chain *Random-2*. Similar cells are randomly distributed along the chain.

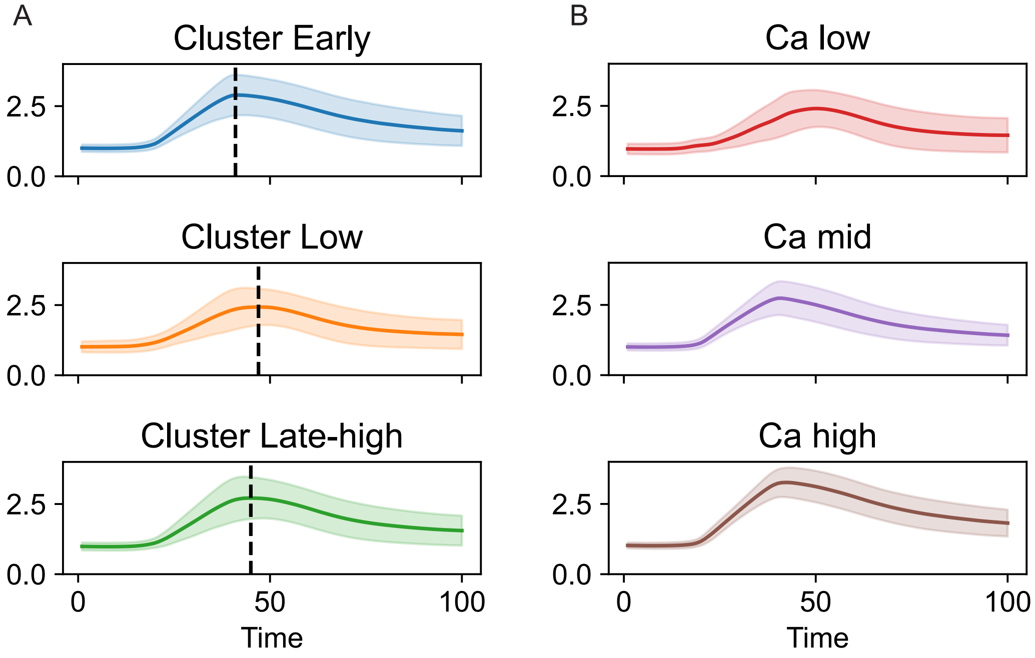


Figure S11: **Comparison of $Ca^{2+}$ dynamics for various cell clusterings of chain *Reduced-3*.** **A**: Simulated $Ca^{2+}$ dynamics sampled from the posterior distributions of cells clustered by posterior parameters. Solid lines mark the mean $Ca^{2+}$ response and shaded region marks one standard deviation. Distinct peak heights and peak times are observed. Dashed lines mark the timing of the $Ca^{2+}$ peak. **B**: Simulated $Ca^{2+}$ dynamics sampled from the posterior distributions of cells clustered by gene expression. Distinct $Ca^{2+}$ peaks are observed (low, mid, high).

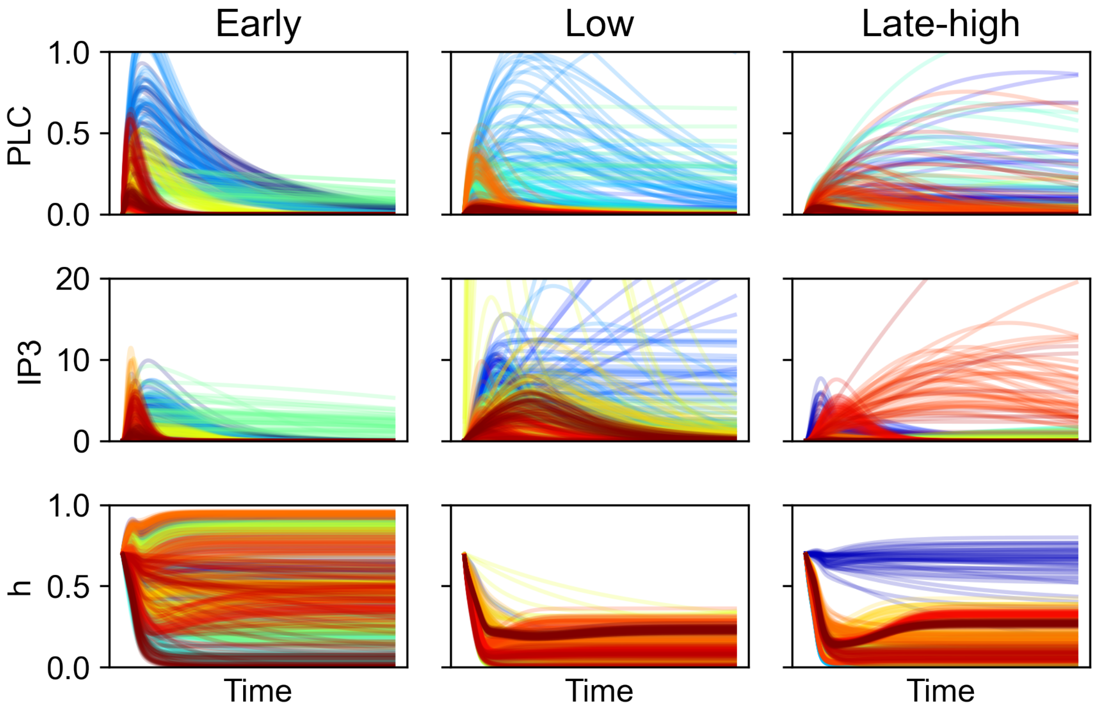


Figure S12: **Comparison of model dynamics in clusters from *Reduced-3* by posterior parameter clustering.** For each of the clusters identified (early, low, and late-high) samples from the posterior distributions of cells belonging to those clusters are plotted for the three model species: PLC, IP3, and $h$. Colors denote samples from different cells. Early responders are characterized by transient peaked trajectories for PLC and IP3. Low responders are characterized by low activation of the IP3 receptor ($h$).

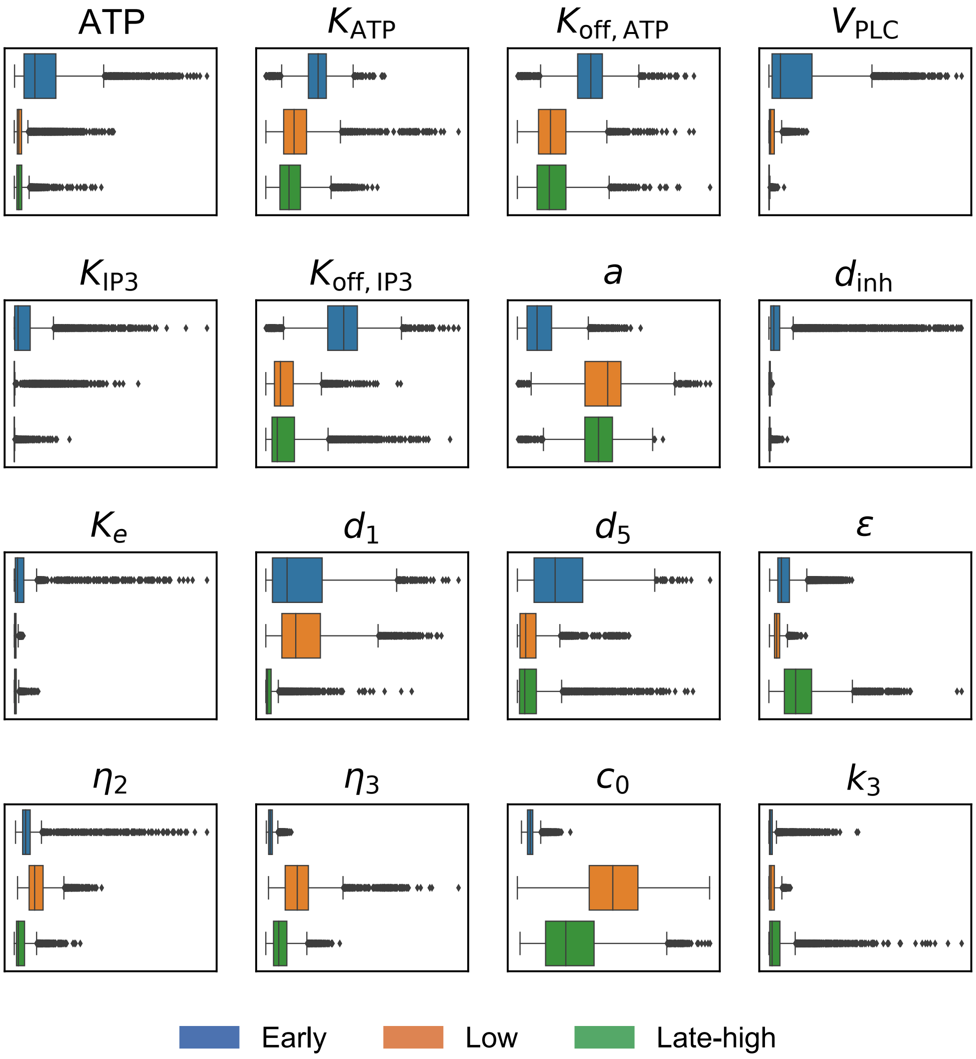


Figure S13: **Comparison of parameter distributions across clusters for *Reduced-3*.** To compare parameter distribution between clusters (for *Reduced-3* clustered by posterior parameters), we performed bootstrap sampling to obtain samples of the same size for each cluster. Boxplots for each parameter in each cluster are shown. We observed different distributions of the bootstrapped samples in each cluster. For instance, low responders had higher concentrations of free Ca$^{2+}$ in ER ($c_0$); early responders differed from others in those parameters controlling the PLC and IP3 dynamics.

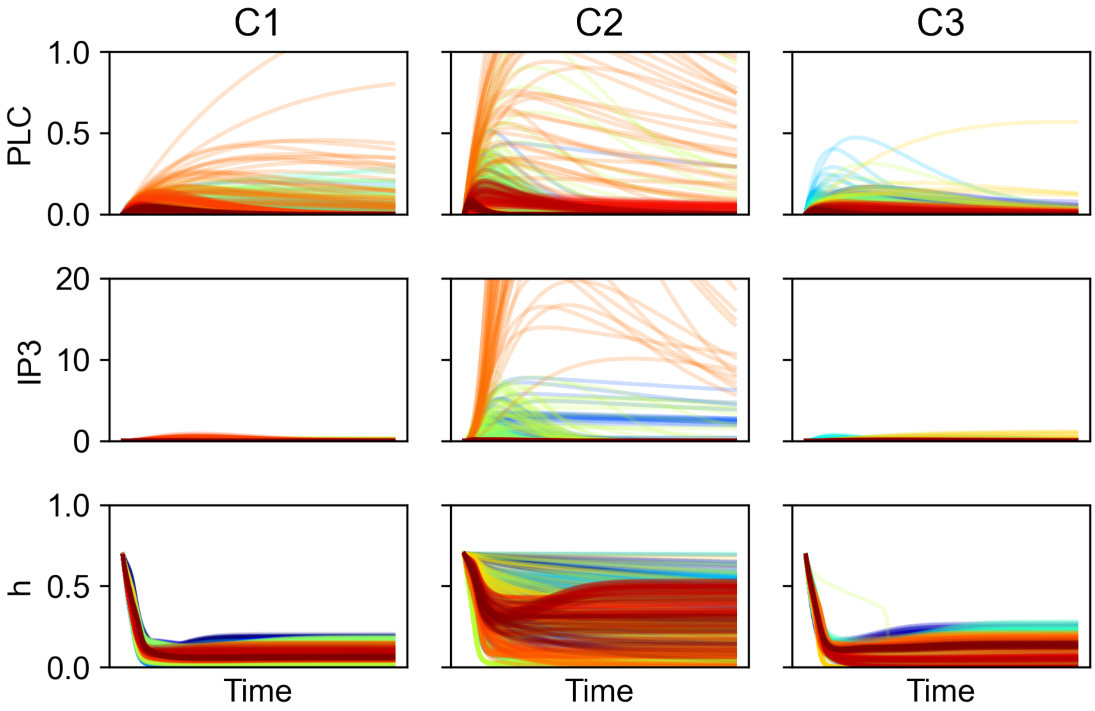


Figure S14: **Comparison of model dynamics in clusters from *Random-2* by posterior parameter clustering.** For each of the clusters identified (C1, C2, C3) samples from the posterior distributions of cells belonging to those clusters are plotted for the three model species: PLC, IP3, and $h$. Colors denote samples from different cells. C2 is characterized by more variable responses than the other clusters; no clearly discernable patterns in the responses are observed.

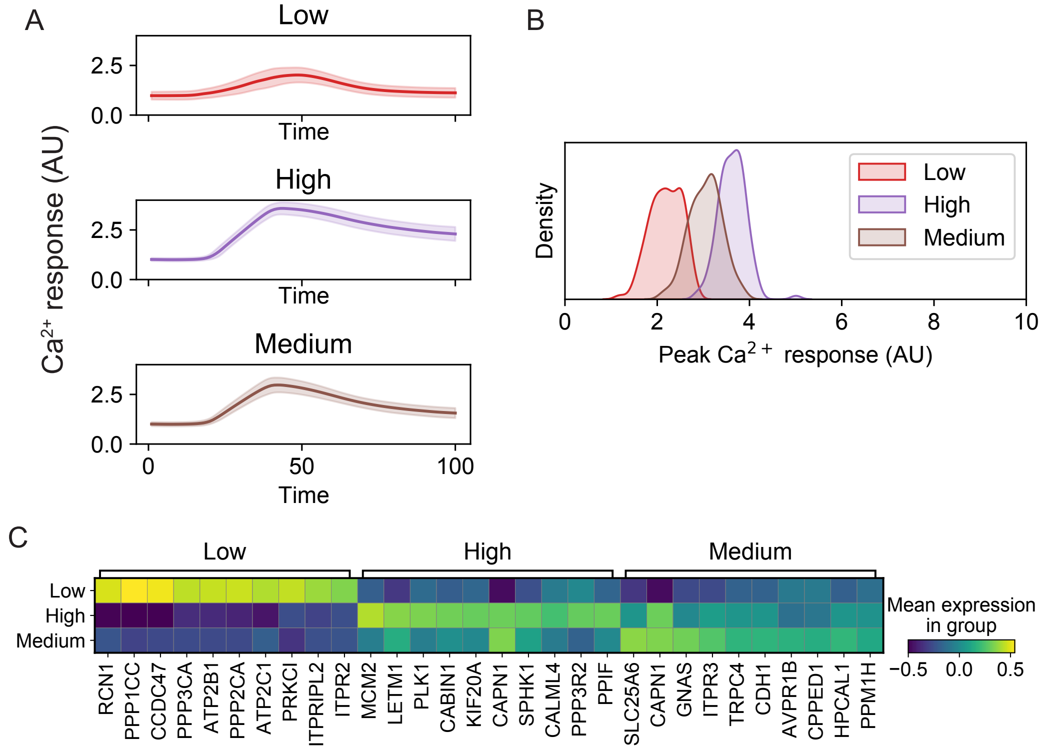


Figure S15: **Clustering cells by $Ca^{2+}$ response profiles.** For 500 cells (same cells as from the chains "*Reduced-3*" and "*Ca-similarity*") we performed $k$-means clustering with $k$=3 on the $Ca^{2+}$ response trajectories of the cells. **A**: $Ca^{2+}$ response trajectories for cells from each of the three clusters identified. The mean $Ca^{2+}$ response (solid line) and the standard deviation (shaded region) are shown. **B**: Kernel density estimation of the distribution of peak $Ca^{2+}$ responses in each cluster. **C**: Marker genes corresponding to each $k$-means cluster.

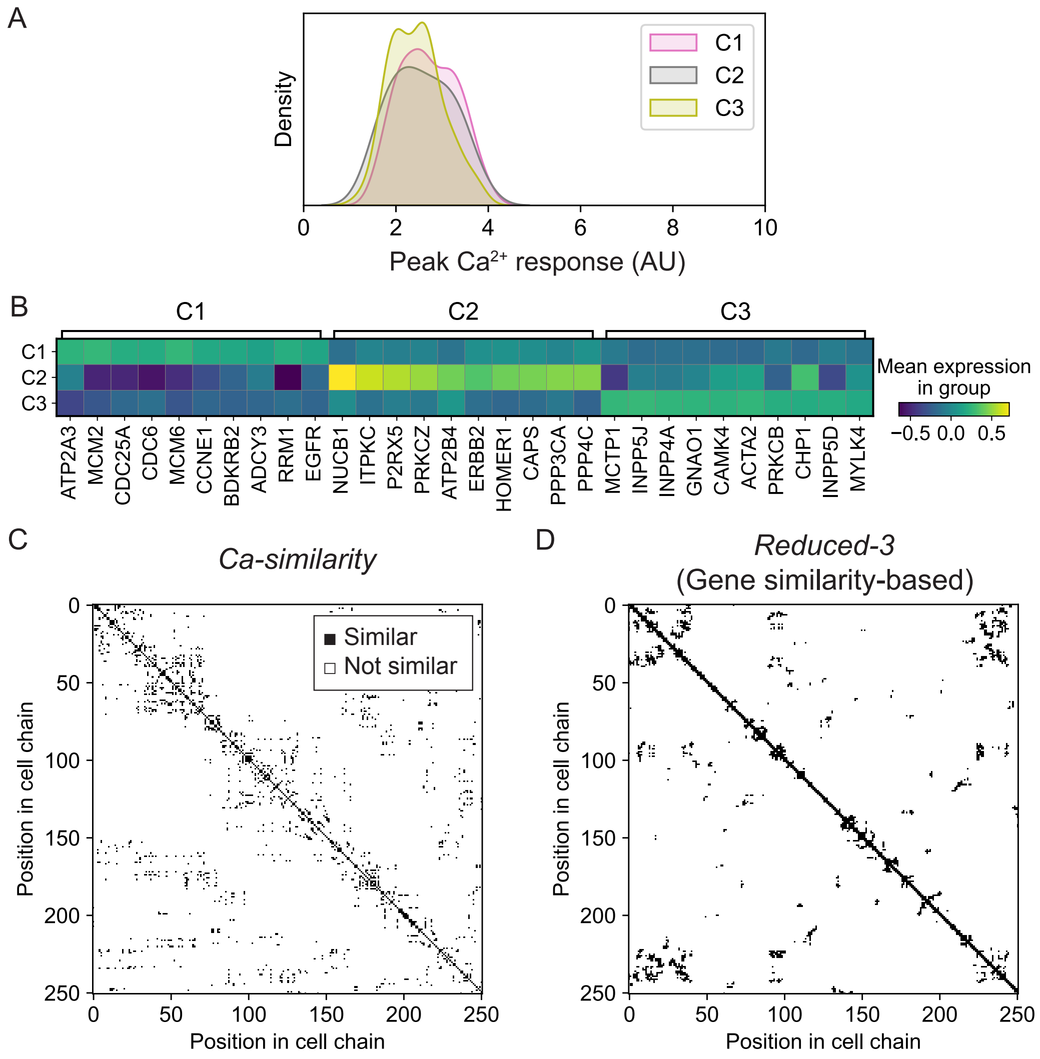


Figure S16: **Posterior parameter clustering for a cell chain constructed via Ca$^{2+}$ signaling similarity.** Inference was performed on cells sampled along a cell chain constructed by Ca$^{2+}$ signaling similarity ("*Ca-similarity*"), followed by hierarchical clustering on the posterior parameter means of each cell. **A**: Kernel density estimation of peak Ca$^{2+}$ response in each posterior parameter cluster. **B**: Top marker genes associated with each cluster. **C**: Gene expression similarity between cells, for cells plotted from "*Ca-similarity*," a cell chain based on similarity in Ca$^{2+}$ signaling. Black pixel indicates that the two corresponding cells are similar in gene expression. **D**: Gene expression similarity between cells, for cells plotted from "*Reduced-3*," a cell chain based on similarity in gene expression.
