## Supplementary Text and Tables for "Single-cell Ca^2+^ parameter inference reveals how transcriptional states inform dynamic cell responses"

**2 Supplementary Tables** **4**

### 1 Supplementary Results

#### 1.1 Analysis of scaling and clipping of prior distributions along cell chains

To assess the use of scaling and clipping the prior standard deviation, we studied to what extent various levels of scaling/clipping control the variance of the posterior distribution. To understand the effects of various choices of scaling/clipping, we ran sampling along the same cell chain using priors derived in three different ways:

1. Use the posterior standard deviation of each parameter from the previous cell as the prior standard deviation for the current cell (*Scale-1.0*)
2. Double the posterior standard deviation from the previous cell (*Scale-2.0*)
3. Scale the posterior standard deviation by 1.5 and clip it to $[0.001, 5]$ (*Similar-r1*)

For *Scale-1.0*, we observed overfitting of $Ca^{2+}$ trajectories. Trajectories simulated from sampled parameters were close to the input trajectories (Figure S2A), but the mean log posterior likelihood kept dropping as more cells were sampled along the cell chain (Figure S2B). Meanwhile, most parameters shrank quickly in standard deviation (Figure S2F–G). For *Scale-2.0*, the marginal distributions of some parameters were stable (Figure S2H), but others grew exponentially in mean and standard deviation (Figure S2I). After the 30th cell, the sampling time also began to increase rapidly (Figure S2C), so we terminated sampling at the 36th cell. *Similar-r1* had stable log posteriors and mean errors (Figure S2A–B). Some parameters still tended to grow or shrink for certain parts of the cell chains, but the clipping technique prevented them from growing or shrinking indefinitely (Figure S2D–E). In other words, using the combination of scaling and clipping, we were able to stabilize parameter distribution along the cell chain. The subsequent cell chain runs in this study adopted these parameters for scaling/clipping.

#### 1.2 Parameter inference for a single cell with multiple epochs

For each cell in Figure 1B–C, we also performed parameter inference with multiple epochs. For the first epoch, we used the Lemon prior. After finishing one epoch, we used the posterior of the epoch to construct the prior for the next epoch as described in Section 2.5. We found that, under the same NUTS configuration, it usually took two to three epochs to reach the same posterior probabilities and fitting errors that were attained by learning from the posterior of a different cell (Figure S4A–F). Improvement in posterior probabilities and fitting errors was usually insignificant after the second or the third epoch (Figure S4A–F), but the marginal posterior distributions of some parameters tended to become narrower in range (Figure S4G–H).

#### 1.3 Parameter inference for a cell chain induced by $Ca^{2+}$ signaling similarity

In addition to sampling along cell chains induced by cell-cell similarity in gene expression, we sampled a cell chain induced by signaling similarity, denoted as *Ca-similarity*. To construct this cell chain from the cells already sampled in *Reduced-3*, we computed the Euclidean distance between the $Ca^{2+}$ response trajectories for each pair of cells, built a graph where each node is a cell and an edge is placed between two cells if their distance is below a threshold, and ran DFS on the graph. The threshold was chosen to be 5.0 so that most cells were still included in the same connected component and the resulting DFS tree was close to a straight chain. We sampled the first 250 cells in *Ca-similarity* and performed parameter inference followed by clustering of cells on their posterior means as with *Reduced-3* and *Random-2* (Section 3.5). The clustering was not able to separate cells by their $Ca^{2+}$ signaling profiles nor their gene expression profiles (Figure S16A–B). A possible reason that the clustering was unable to divide cells into distinct populations is that the *Ca-similarity* cell chain did not permit a transfer of transcriptional information from cell to cell. We observed that most cells in *Ca-similarity* were not similar in gene expression to their immediate neighbors – predecessor or successor – even though we do observe structure (similarity blocks) in the heatmap (Figure S16C). In constrast, every cell in *Reduced-3* was similar in gene expression with both of its immediate neighbors (Figure S16D). Thus, in the case of *Ca-similarity*, information encoded within parameter posteriors regarding transcriptional states might have been lost as we sampled cells along the cell chain.

### 2 Supplementary Tables

| Run name | Sampled in chain? | Similarity-based? | Reduced model | NUTS hyperparameters (`warmup`, `max_tree_depth`) |
|---|---|---|---|---|
| *Indiv. cells run 1* | N | – | N | (500, 15) |
| *Indiv. cells run 2* | N | – | N | (1000, 15) |
| *Similar-r1* | Y | Expression-based | N | (500, 10) |
| *Similar-r2* | Y | Expression-based | N | (1000, 15) |
| *Similar-r3* | Y | Expression-based | N | (500, 15) |
| *Reduced-1* | Y | Expression-based | Constant $\eta_1$ | (500, 10) |
| *Reduced-2* | Y | Expression-based | Constant $B_e$ | (500, 10) |
| *Reduced-3* | Y | Expression-based | Constant $\eta_1$ and $B_e$ | (500, 10) |
| *Random-1* | Y | N | Constant $\eta_1$ and $B_e$ | (500, 10) |
| *Random-2* | Y | N | Constant $\eta_1$ and $B_e$ | (500, 10) |
| *Ca-similarity* | Y | $Ca^{2+}$-based | Constant $\eta_1$ and $B_e$ | (500, 10) |

Table S1: Summary of inference runs referred to in this work. “Sampled in chain” refers to whether the run is cell predecessor-based (cell prior informed by the previous cell’s posterior). “Similarity-based” refers to whether the cell chain ordering is informed by cell-cell similarity in either gene expression or $Ca^{2+}$ signaling.

| Cell ID | *Similar-r1* | *Indiv. cells run 1* | *Indiv. cells run 2* |
|---|---|---|---|
| 5121 | $441.07 \pm 17.23$ | $449.61 \pm 3.26$ | $449.55 \pm 3.39$ |
| 5125 | $440.46 \pm 4.43$ | $337.23 \pm 21.41$ | $326.50 \pm 3.52$ |
| 5116 | $398.47 \pm 3.45$ | $344.46 \pm 3.39$ | $343.90 \pm 3.54$ |
| 5107 | $378.01 \pm 21.79$ | $322.17 \pm 58.75$ | $337.87 \pm 69.66$ |
| 5093 | $382.29 \pm 3.39$ | $362.15 \pm 2.97$ | $358.37 \pm 7.01$ |
| 5127 | $408.79 \pm 3.48$ | $302.19 \pm 68.33$ | $365.09 \pm 14.78$ |
| 5117 | $408.77 \pm 3.33$ | $363.86 \pm 9.18$ | $359.55 \pm 8.14$ |
| 5046 | $385.26 \pm 3.16$ | $218.81 \pm 7.49$ | $224.66 \pm 3.34$ |
| 5091 | $356.95 \pm 3.33$ | $311.55 \pm 17.02$ | $322.25 \pm 10.83$ |
| 5082 | $329.24 \pm 3.29$ | $322.96 \pm 4.15$ | $323.99 \pm 3.84$ |

Table S2: Comparison of mean log posterior for ten cells from the similarity-based chain (*Similar-r1*) with the same cells sampled individually using uninformative priors with either 500 or 1000 warmup steps (*Indiv. cells run 1* and *Indiv. cells run 2*). For each fitted cell the log posterior is given $\pm$ the standard deviation.

| Run | Maximum tree depth | Number of warmup steps | Number of converged cells | Number of mixed chains | $\widehat{R}$ (mixed only) |
|---|---|---|---|---|---|
| *Indiv. cells run 1* | 15 | 500 | 10 | $3.80 \pm 0.40$ | $1.77 \pm 0.79$ |
| *Indiv. cells run 2* | 15 | 1000 | 9 | $3.89 \pm 0.31$ | $1.70 \pm 0.86$ |
| *Similar-r1* | 10 | 500 | 10 | $3.80 \pm 0.40$ | $1.04 \pm 0.10$ |

Table S3: Sampling performance of the similarity-based chain (*Similar-r1*) and two runs in which the same cells were sampled individually using the uninformative Lemon prior with either 500 or 1000 warmup steps (*Indiv. cell run 1* and *Indiv. cell run 2*). The number of mixed chains (out of 4) and the average $\widehat{R}$ are used to compare performance, each is given $\pm$ the standard deviation.

| Run | Sampling time (minutes) | Mean error |
|---|---|---|
| *Reduced-3* | $67.25 \pm 83.62$ | $0.79 \pm 0.37$ |
| *Random-1* | $64.84 \pm 46.43$ | $0.82 \pm 0.47$ |

Table S4: Comparison between cell chains sampled using similarity-informed priors (*Reduced-3*) or randomly ordered (*Random-1*). The mean sampling times and mean errors between fitted trajectories and data are given $\pm$ the standard deviation. No significant differences are observed between the two chains. Both cell chains use a reduced model in which $B_e$ and $\eta_1$ were set to constant values.

| Run | Number of warmup steps | Max tree depth | Tree depth | Sampling time (minutes) | $\widehat{R}$ | Mean error |
|---|---|---|---|---|---|---|
| *Similar-r1* | 500 | 10 | $9.76 \pm 0.43$ | $68.11 \pm 38.75$ | $1.11 \pm 0.33$ | $0.72 \pm 0.32$ |
| *Similar-r2* | 1000 | 15 | $10.36 \pm 0.80$ | $253.23 \pm 151.75$ | $1.11 \pm 0.40$ | $0.65 \pm 0.29$ |
| *Similar-r3* | 500 | 15 | $10.72 \pm 1.21$ | $189.48 \pm 154.54$ | $1.08 \pm 0.36$ | $0.66 \pm 0.29$ |

Table S5: Comparison between cell chains run with different hyperparameters, all based on similarity-informed cell chains. The tree depth reported is the mean over the population of 500 fitted cells. All statistics are given $\pm$ the standard deviation.

| Run | Tree depth | Sampling time (minutes) | $\widehat{R}$ | Mean error |
|---|---|---|---|---|
| *Similar-r1* | $9.76 \pm 0.44$ | $55.54 \pm 34.67$ | $1.12 \pm 0.37$ | $0.79 \pm 0.37$ |
| *Reduced-1* | $9.61 \pm 0.66$ | $52.73 \pm 39.89$ | $1.14 \pm 0.41$ | $0.84 \pm 0.42$ |
| *Reduced-2* | $9.61 \pm 0.74$ | $58.27 \pm 66.76$ | $1.18 \pm 0.50$ | $0.84 \pm 0.55$ |
| *Reduced-3* | $9.86 \pm 0.35$ | $67.25 \pm 83.62$ | $1.17 \pm 0.47$ | $0.79 \pm 0.37$ |

Table S6: Comparison between cell chains run on the full model (*Similar-r1*) and models reduced by setting one or two parameters to a constant. All statistics are given $\pm$ the standard deviation. No significant differences are observed between the chains.

| Parameter | Mean | Standard deviation | Mean sensitivity (0.01-quantile) | Mean sensitivity (0.99-quantile) |
|---|---|---|---|---|
| ATP | 0.007 | 0.010 | 8.98 | 14.0 |
| $K_{\text{ATP}}$ | 0.032 | 0.017 | 5.43 | 4.47 |
| $K_{\text{off, ATP}}$ | 0.031 | 0.016 | 5.57 | 4.50 |
| $V_{\text{PLC}}$ | 1.46 | 3.75 | 8.93 | 13.5 |
| $K_{\text{IP3}}$ | 0.207 | 0.548 | 17.3 | 6.86 |
| $K_{\text{off, IP3}}$ | 0.043 | 0.038 | 4.31 | 3.52 |
| $a$ | 0.017 | 0.008 | 2.87 | 1.78 |
| $d_{\text{inh}}$ | 0.810 | 2.80 | 1.28 | 2.01 |
| $K_e$ | 0.005 | 0.014 | 0.47 | 0.60 |
| $d_1$ | 2.62 | 3.50 | 25.0 | 7.40 |
| $d_5$ | 0.870 | 1.01 | 2.64 | 2.68 |
| $\epsilon$ | 0.347 | 0.283 | 13.3 | 49.5 |
| $\eta_2$ | 0.223 | 0.224 | 13.8 | 58.2 |
| $\eta_3$ | 0.726 | 0.667 | 32.2 | 9.77 |
| $c_0$ | 20.0 | 14.2 | 9.65 | 38.1 |
| $k_3$ | 0.075 | 0.140 | 1.31 | 1.26 |

Table S7: Statistics of the posterior distribution for cells fit to the similarity-based chain and mean parameter sensitivities, reduced model (*Reduced-3*). First, marginal posterior means of each parameter are computed for each cell, we then compute the mean and standard deviation of the posterior means of each parameter over all cells, i.e. across the cell population. Mean sensitivities are taken from Figure 3B.

| Gene | Parameter | Correlation | p-value |
|---|---|---|---|
| PPP1CC | $\eta_3$ | $0.382423$ | $1.752063 \times 10^{-15}$ |
| CCDC47 | $\eta_3$ | $0.373109$ | $9.304525 \times 10^{-15}$ |
| PPP1CC | $a$ | $0.355153$ | $2.005634 \times 10^{-13}$ |
| MCM6 | $K_{\text{off, IP3}}$ | $-0.343088$ | $1.419453 \times 10^{-12}$ |
| PPP2CA | $\eta_3$ | $0.337860$ | $3.229559 \times 10^{-12}$ |
| ITPRIPL2 | $\eta_3$ | $0.335172$ | $4.899232 \times 10^{-12}$ |
| RCN1 | $\eta_3$ | $0.333198$ | $6.636243 \times 10^{-12}$ |
| MCM6 | $c_0$ | $0.329770$ | $1.118234 \times 10^{-11}$ |
| MCM6 | $\eta_3$ | $0.326605$ | $1.799954 \times 10^{-11}$ |
| CCDC47 | $K_{\text{off, IP3}}$ | $-0.324868$ | $2.332030 \times 10^{-11}$ |
| PPP1CC | $K_{\text{off, IP3}}$ | $-0.319847$ | $4.883968 \times 10^{-11}$ |
| PPP3CA | $\eta_3$ | $0.319310$ | $5.281702 \times 10^{-11}$ |
| RRM1 | $K_{\text{off, IP3}}$ | $-0.318196$ | $6.209595 \times 10^{-11}$ |
| PRKCI | $\eta_3$ | $0.308928$ | $2.325989 \times 10^{-10}$ |
| PIP4K2C | $\eta_3$ | $0.308875$ | $2.343277 \times 10^{-10}$ |
| PPP1CC | $c_0$ | $0.308291$ | $2.542612 \times 10^{-10}$ |
| RRM1 | $\eta_3$ | $0.303851$ | $4.703189 \times 10^{-10}$ |
| E2F1 | $K_{\text{off, IP3}}$ | $-0.301569$ | $6.426301 \times 10^{-10}$ |
| CCNE1 | $K_{\text{off, IP3}}$ | $-0.292604$ | $2.133973 \times 10^{-9}$ |
| CCNE1 | $c_0$ | $0.284548$ | $6.057618 \times 10^{-9}$ |

Table S8: Gene-parameter pairs ranked by correlation coefficient in a similarity-based chain (*Reduced-3*). Top variable genes were determined (see Section 2.7) and for all possible pairs of variable genes and model parameters we computed the Pearson correlation between the gene expression values and the posterior parameter means for all cells in the population. Then the gene-parameter pairs were ranked by their absolute Pearson correlation coefficients (top 20 pairs listed).

| Gene | Parameter | Correlation | p-value |
| --- | --- | --- | --- |
| SLC25A6 | $K_{\text{ATP}}$ | $0.200663758$ | $3.11 \times 10^{-19}$ |
| SLC25A6 | $K_{\text{off, ATP}}$ | $0.180954575$ | $7.11 \times 10^{-16}$ |
| CCDC47 | $d_5$ | $0.17514209$ | $5.93 \times 10^{-15}$ |
| ATP2B1 | $d_5$ | $0.165159467$ | $1.92 \times 10^{-13}$ |
| PPP1CC | $d_5$ | $0.155976354$ | $3.92 \times 10^{-12}$ |
| PRKCI | $d_5$ | $0.150758764$ | $2.01 \times 10^{-11}$ |
| RCN1 | $\eta_3$ | $0.150008815$ | $2.54 \times 10^{-11}$ |
| RCN1 | $d_5$ | $0.149302892$ | $3.15 \times 10^{-11}$ |
| ATP2B1 | $\eta_3$ | $0.14738222$ | $5.63 \times 10^{-11}$ |
| PPP1CC | $\eta_3$ | $0.144494084$ | $1.33 \times 10^{-10}$ |
| PPP2CA | $a$ | $0.144203999$ | $1.45 \times 10^{-10}$ |
| CCDC47 | $a$ | $0.143404138$ | $1.84 \times 10^{-10}$ |
| PPP3CA | $d_5$ | $0.143021068$ | $2.05 \times 10^{-10}$ |
| PPM1F | $d_5$ | $0.141377079$ | $3.31 \times 10^{-10}$ |
| PPP3CA | $\eta_3$ | $0.139341446$ | $5.94 \times 10^{-10}$ |
| PPP1CC | $a$ | $0.137844169$ | $9.08 \times 10^{-10}$ |
| RCN1 | $a$ | $0.137339719$ | $1.05 \times 10^{-9}$ |
| CCDC47 | $k_3$ | $0.137063766$ | $1.13 \times 10^{-9}$ |
| ATP2C1 | $d_5$ | $0.136528503$ | $1.31 \times 10^{-9}$ |
| SLC25A6 | $d_5$ | $-0.136192465$ | $1.44 \times 10^{-9}$ |

Table S9: Gene-parameter pairs ranked by correlation coefficient in a randomly ordered chain (*Random-2*). Top variable genes were determined (see Section 2.7) and for all possible pairs of variable genes and model parameters we computed the Pearson correlation between the gene expression values and the posterior parameter means for all cells in the population. Then the gene-parameter pairs were ranked by their absolute Pearson correlation coefficients (top 20 pairs listed).
